# Determinants of ligand-functionalized DNA nanostructure-cell interactions

**DOI:** 10.1101/2021.02.24.432702

**Authors:** Glenn A.O. Cremers, Bas J.H.M. Rosier, Ab Meijs, Nicholas B. Tito, Sander M.J. van Duijnhoven, Hans van Eenennaam, Lorenzo Albertazzi, Tom F.A. de Greef

**Affiliations:** Laboratory of Chemical Biology and Institute for Complex Molecular Systems, Eindhoven University of Technology, P.O. Box 513, 5600 MB Eindhoven, The Netherlands; Computational Biology Group, Department of Biomedical Engineering, Eindhoven University of Technology, P.O. Box 513, 5600 MB Eindhoven, The Netherlands; Department of Biosystems Science and Engineering, ETH Zurich, Mattenstrasse 26, 4058 Basel, Switzerland; Molecular Biosensing for Medical Diagnostics, Department of Biomedical Engineering, Eindhoven University of Technology, P.O. Box 513, 5600 MB Eindhoven, The Netherlands; Aduro Biotech Europe B.V., Oss, the Netherlands; Electric Ant Lab, Science Park 106, 1098 XG, Amsterdam, The Netherlands; Institute for Molecules and Materials, Radboud University, Heyendaalseweg 135, 6525 AJ Nijmegen, The Netherlands

## Abstract

Synthesis of ligand-functionalized nanomaterials with control over size, shape and ligand orientation, facilitates the design of tailored nanomedicines for therapeutic purposes. DNA nanotechnology has emerged as a powerful tool to rationally construct two- and three-dimensional nanostructures, enabling site-specific incorporation of protein ligands with control over stoichiometry and orientation. To efficiently target cell surface receptors, exploration of the parameters that modulate cellular accessibility of these nanostructures is essential. In this study we systematically investigate tunable design parameters of antibody-functionalized DNA nanostructures binding to therapeutically relevant receptors. We show that, although the native affinity of antibody-functionalized DNA nanostructures remains unaltered, the absolute number of bound surface receptors is lower compared to soluble antibodies and is mainly governed by nanostructure size and DNA handle location. The obtained results provide key insights in the ability of ligand-functionalized DNA nanostructures to bind surface receptors and yields design rules for optimal cellular targeting.

## Introduction

In the last decades, nanoscale materials have emerged as a promising biomedical tool for diagnosis and treatment of diseases.^1–3^ Nanomedicines are a class of nanomaterials which can be constructed from polymeric, inorganic, or organic particles containing biologically active ligands, and are specifically formulated to induce cellular signaling mediated by ligand-receptor binding or to deliver therapeutic drugs to specific cells or tissues.^4,5^ Incorporation of multiple ligands onto nanoparticles results in a higher avidity towards target receptors as a result of multivalency ^6,7^, and facilitates local delivery which increases drug accumulation in the site of interest, enhancing therapeutic efficiency and reducing off-target effects. Optimization of the synthesis and formulation of nanomedicines has revealed several parameters that modulate targeting efficiency and cellular uptake^8^, which include the orientation^9^, mobility^10^, and surface density of ligands on the nanoparticle.^11–13^ In addition, nanoparticle size, shape, and aspect ratio also influence their uptake and therapeutic effectiveness.^14–17^ For example, rod-shaped nanoparticles display more efficient cell binding compared to spherical nanoparticles,^18^ whereas spherical particles more efficiently enhance cellular uptake.^19^ To further unlock the potential of nanomedicines it is crucial to control the synthesis of the nanoscale vehicles and, as such, elucidate critical design parameters for cellular targeting as a function of vehicle composition, shape, size, and geometry.

The programmability of DNA origami can be employed to construct well-defined nanostructures that allow site-specific immobilization of ligands with unprecedented control over stoichiometry and orientation.^20,21^ DNA nanostructures have been used as delivery vehicles by selectively encapsulating drug molecules that can be released in a controlled fashion when the DNA nanostructure binds to specific cell types.^22,23^ Additionally, these nanostructures can be used to study distance effects of receptor activation with nanometer precision^24–28^, enhance the cellular uptake of therapeutic drugs^29,30^, and are able to modulate drug release kinetics.^31,32^ More specifically, it has been shown that compact nanostructures with a low aspect ratio are the preferred delivery vehicles for internalization^33^ and that larger DNA origami structures exhibit a higher uptake efficiency.^34^ Some of the initial challenges for the use of DNA nanostructures for biomedical applications have been addressed and overcome, including low-scale inefficient production, poor structural integrity in physiological fluids and degradation by nuclease activity, making DNA origami-based nanostructures a potential platform for the design of tailored nanomedicines.^35–42^

To maximize the potential of DNA nanostructures as a generic platform for precision medicine it is essential to analyze all parameters that influence nanostructure performance. While the parameters that modulate cellular uptake are relatively well understood, it is currently unclear if DNA nanostructures interfere with the interaction between ligands and cellular surface receptors. Although research has shown that incorporation of a protein ligand onto a DNA nanostructure does not alter the native affinity of the ligand for the receptor^24,43^, the crowded and irregularly shaped cell surface could interfere with binding of ligand-functionalized DNA nanostructures to surface receptors as a result of steric hindrance. This can lead to ineffective cellular binding of DNA nanostructure-based nanomedicines, and subsequently to decreased downstream signaling efficiency and reduced therapeutic effectiveness.

In this study, we aim to systematically evaluate key parameters that modulate surface receptor binding of antibody-functionalized DNA nanostructures (Fig. **1a**). As a model platform, we investigate receptor binding to multiple cellular surface receptors, including PD-1, EGFR and HER-2, using 18-helix bundle DNA nanorods functionalized with a single antibody.^24,43^ We employ stoichiometric fluorescent labeling of antibodies to quantitatively assess cellular accessibility and show that the DNA nanorod limits the absolute number of cellular surface receptors that are bound compared to the corresponding free antibody, although the native affinity of the antibody remains unaltered. Subsequently, we use the cancer immunotherapy-related PD1 receptor^44^ to study individual determinants that govern receptor accessibility in more detail. Taking advantage of the spatial addressability of DNA origami, we provide direct evidence that DNA handle location, in contrast to linker length and electrostatic interactions, is a key parameter for optimal receptor binding. We then design multiple DNA origami structures and observe a negative correlation between receptor targeting efficiency and DNA nanostructure size. To understand the role of cellular determinants, we demonstrate that receptor accessibility is also influenced by surface receptor density and the presence of glycoproteins. Finally, we show that limited cellular accessibility of anti-PD1-functionalized DNA nanorods results in ineffective blocking of cellular PD1/PDL1 interactions *in vitro*. Taken together, our analysis provides key insights on the parameters that modulate receptor accessibility and can be used to guide the design of DNA origami nanostructures for optimal cellular targeting.

**Figure 1.**
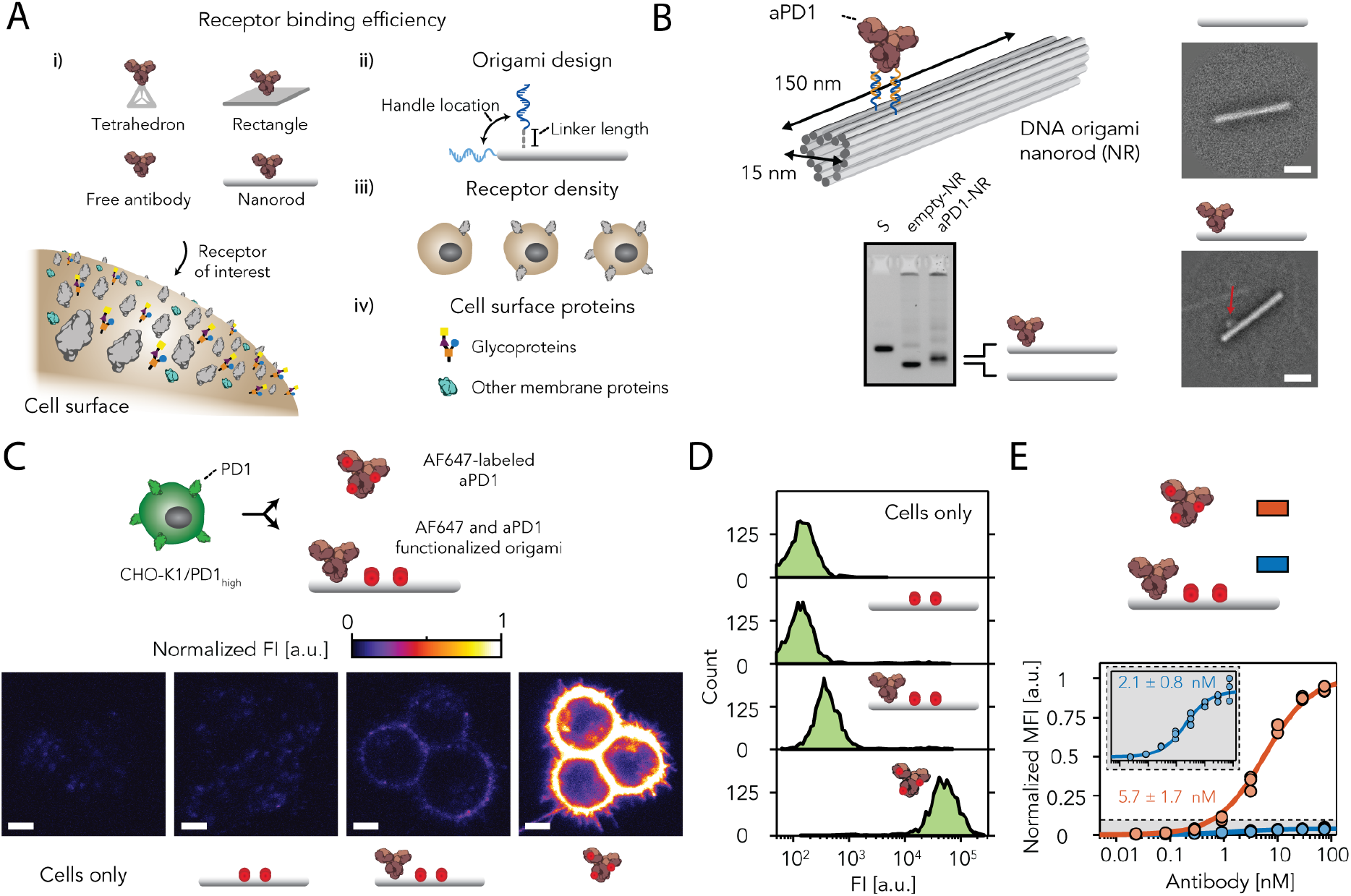
Targeting cellular surface receptors using antibody-functionalized DNA nanorods. **(a)** Schematic overview of the determinants that potentially modulate receptor binding of antibody-functionalized DNA nanostructures, including (i) DNA nanostructure size and shape, (ii) tunable DNA origami design parameters, (ii) receptor density and (iii) the presence of other cell surface proteins. **(b)** Reference-free class averages obtained from single-particle TEM micrographs and electrophoretic mobility analysis of a self-assembly reaction of the 18-helix bundle nanorod with-(aPD1-NR) and without (empty-NR) the site-specific incorporation of an anti-programmed cell death protein 1 antibody (aPD1). Scale bar, 50 nm, labels: S, scaffold **(c)** Confocal images and **(d)** flow cytometric analysis of PD1-overexpressing Chinese hamster ovary K1 (CHO-K1_PD1-high_) cells incubated for 1h with 20 nM of Alexa-647-labeled (AF647) empty-NR, aPD1-NR or free aPD1. Scale bar, 5 μm. **(e)** Flow cytometric analysis of AF647-labeled aPD1 and aPD1-NR titrated to CHO-K1_PD1-high_ cells for 1h. The mean fluorescent intensities, corrected for AF67 labeling efficiency, were fitted to a non-cooperative Hill equation and normalized to the fitted maximum fluorescence intensity of cells stained with AF647-aPD1 to extract the apparent dissociation constant. Individual data points represent the normalized mean fluorescent intensity (MFI) of 2000 gated single-cell events (n = 3 technical replicates).

## Results

### Targeting cellular surface receptors with antibody-functionalized DNA nanorods

To investigate the role of DNA nanostructures in the interaction between ligands and cellular surface receptors we constructed an 18-helix bundle DNA nanorod^24,31^ (NR, 15×150 nm^2^) functionalized with a single anti-PD1 antibody. We previously developed a modular conjugation strategy to site-selectively couple ssDNA handles to the Fc domain of antibodies using a small photocrosslinkable protein G adaptor that ensures correct antibody orientation on the DNA nanostructure.^43^ In this work we employed this method to site-specifically conjugate two ssDNA anti-handles to the Fc region of anti-PD1 antibodies (aPD1) that hybridizes to two complementary ssDNA handles protruding from the NR surface (**Supplementary Figs. 1-3**). Agarose gel electrophoresis confirmed the self-assembly of NRs and transmission electron microscopy revealed site-specific incorporation of DNA-aPD1 conjugates on the NR surface (**Fig. 1b** and **Supplementary Figs. 4** and **5**). Next, we used an engineered Chinese hamster ovary (CHO-K1) cell line stably expressing a high level of PD1 receptors to analyze receptor binding efficiency of AF647-labeled aPD1-NR and compare it to AF647-labeled free aPD1. We confirmed that the structural integrity of NRs was maintained during cellular labeling while confocal microscopy analysis revealed that both aPD1-NR and free aPD1 are localized on the cellular membrane demonstrating successful binding (**Fig. 1c** and **Supplementary Fig. 6**). In addition, flow cytometric analysis of CHO-K1_PD1-high_ cells showed an increase in mean fluorescence intensity of individual cells labeled with either aPD1-NR or free aPD1 (**Fig. 1d**). In both measurements, however, the absolute fluorescent intensity of CHO-K1_PD1-high_ cells incubated with aPD1-NR was approximately 10-fold lower than cells incubated with free aPD1. To exclude the possibility that the difference in fluorescent intensity is due to enhanced internalization of free aPD1 compared to origami-tethered aPD1, we treated aPD1-labeled cells with an acidic solution to remove cell surface aPD1.^45^ Subsequently, we measured fluorescent intensity levels using flow cytometry to determine internalized fluorescent signals. The relative decrease in fluorescent intensity levels after acid treatment was similar for CHO-K1_PD1-high_ cells treated with free AF647-aPD1 or AF647-labeled aPD1-NR, demonstrating that aPD1 internalization is not the primary contributor to the difference in fluorescent intensity (**Supplementary Fig. 7**). We also tested whether the low fluorescent signal might be a result of the purification method used to remove uncoupled aPD1-DNA conjugates from functionalized aPD1-NR nanostructures. Purification of aPD1-NRs using agarose gel extraction^46^ instead of two rounds of PEG precipitation, however, showed similar differences in fluorescent intensity between free AF647-aPD1 and AF647-labeled aPD1-NRs, excluding the purification method as a main source for the low levels of fluorescent intensity (**Supplementary Fig. 8**). Collectively, these results indicate that NRs limit PD1 binding.

To assess the impact of NRs on receptor binding affinity, we titrated AF647-labeled aPD1-NR or free aPD1 against CHO-K1_PD1-high_ cells and measured the mean fluorescence intensity using flow cytometry.^43,47^ After correcting for AF647 labeling efficiency (**Supplementary Fig. 9**), CHO-K1_PD1-high_ labeling with free aPD1 resulted in an absolute fluorescent intensity 20-fold higher compared to aPD1-NR labeling (**Fig. 1e**). Surprisingly, this large difference in fluorescent intensity did not translate to a different apparent dissociation constant (K_D,app_) of free aPD1 and aPD1-NR, indicating that aPD1 retained its affinity when immobilized onto NRs. These experimental results were rationalized using a thermodynamic model that describes binding of antibodies to surface-tethered receptors and is able to explain the observed difference in fluorescent intensity between free aPD1 and aPD1-NR in relation to the measured the K_D,app_ (**Supplementary Notes** and **Supplementary Fig. 26**). Using the model, we show that K_D,app_, in contrast to the absolute fluorescent intensity, is independent of the absolute number of bound receptor binding sites and only a function of the fractional occupancy of cell surface receptors. Experimentally this was verified by titrating AF647-labeled aPD1 to CHO-K1 cells expressing low, intermediate, and high levels of PD1, respectively (**Supplementary Fig. 10**). Translating these results to the experimental data of aPD1-NR binding to CHO-K1_PD1-high_ cells (**Fig. 1e**) we therefore hypothesized that steric hindrance of NRs limits the absolute number of receptors that can bind to aPD1-NR nanostructures. Taken together, our results reveal that the native affinity of DNA origami-tethered aPD1 antibodies remains unaltered compared to free aPD1 but that the absolute number of bound DNA nanostructures is lower compared to the free antibody, resulting in a lowered binding efficiency.

### Quantifying availability of cellular surface receptors to DNA nanorods

Having shown that the DNA nanorod limits receptor binding efficiency of DNA origami-tethered aPD1 antibodies to CHO-K1_PD1-high_, we sought to quantify the absolute availability of cellular receptors to DNA nanostructures. We therefore developed a general assay, using a two-step labeling method, in which we first labeled cells with DNA-antibody conjugates and subsequently incubated DNA-antibody-labeled cells with either a small CY5-functionalized imager strand (CY5-IM) or a CY5-functionalized DNA nanorod (CY5-NR) (**Fig. 2a**). DNA-antibody conjugates are site-specifically functionalized with two ssDNA handles and are therefore either available to two fluorescently labeled imager strands or a single CY5-NR (**Fig. 1b**). To establish proof of concept, we titrated DNA-aPD1 conjugates to CHO-K1_PD1-high_ and fluorescently labeled the cells using a fixed concentration of imager or NR and obtained similar K_D,app_ as previously determined (compare **Fig. 1e and 2b**), while the absolute fluorescent intensities showed over 10-fold difference. Next, we evaluated receptor binding efficiency of NRs to A431 and SKBR3 cells, expressing therapeutically relevant epidermal growth factor receptors (EGFR) and human epidermal growth factor receptors 2 (HER-2), respectively. To this end therapeutic antibodies Cetuximab (anti-EGFR) and Trastuzumab (anti-HER2) were conjugated to two ssDNA handles and titrated to A431 and SKBR3 cells, respectively (**Supplementary Fig. 11**). Both experimental results similarly confirmed that receptor binding efficiency was decreased when cells were labeled with NRs, while K_D,app_ remained unaltered (**Fig. 2b**).

**Figure 2.**
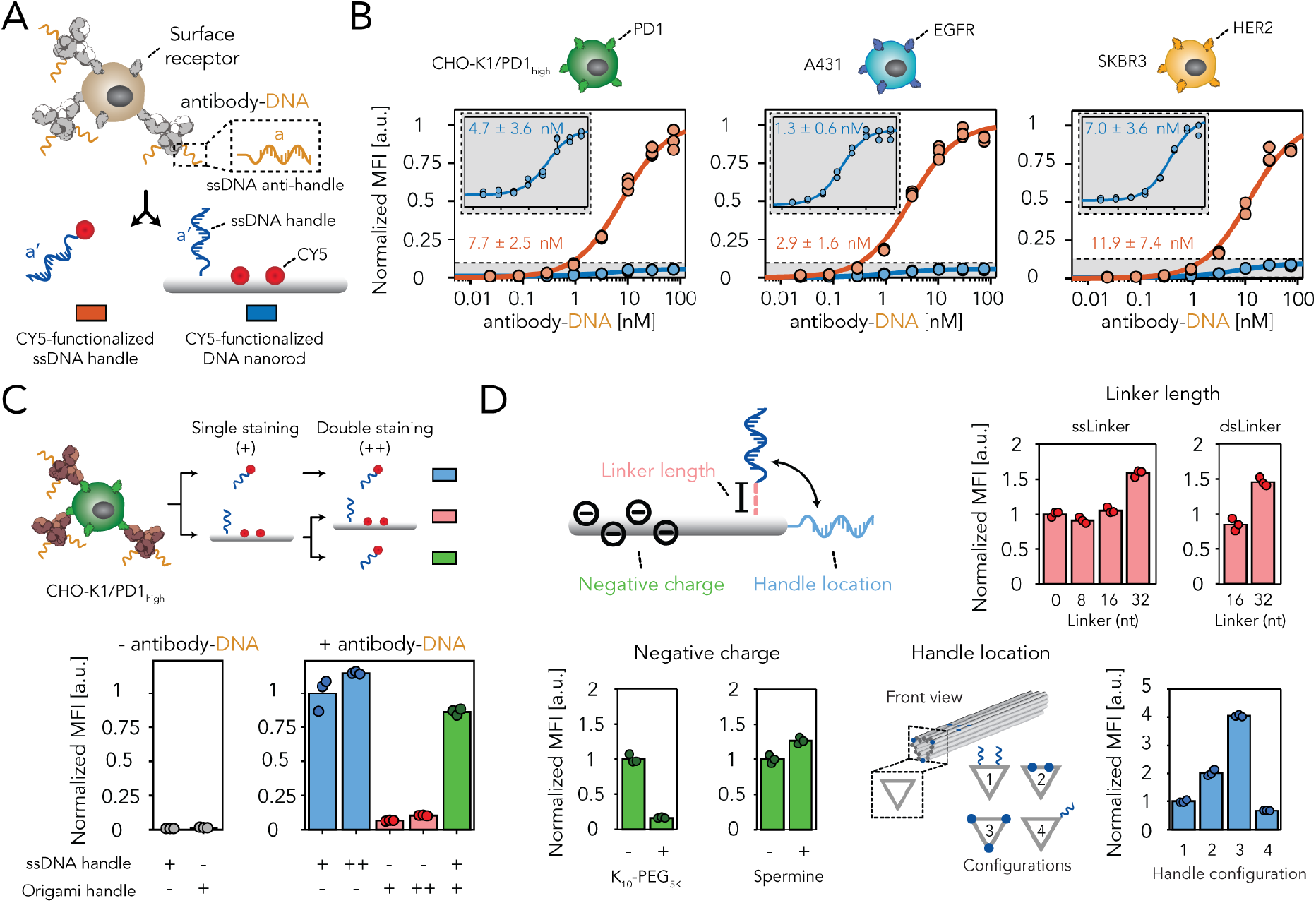
Quantification of DNA nanostructure-cell surface receptor interactions. **(a)** Schematic of the experimental set-up to quantitively assess the absolute fraction of surface receptors targeted using DNA nanostructures. 20 nM of antibodies, site-specifically labeled with 2 DNA handles, were incubated for 30 min with cells expressing target receptors and subsequently labeled with 10 nM of a complementary CY5-labeled imager strand or a CY5-functionalized DNA nanorod that includes complementary handle-extended staple strands for 30 min. **(b)** Flow cytometric analysis of DNA nanostructure receptor targeting in three cell lines (CHO-K1_PD1-high_, A431 and SKBR3) expressing the programmed cell death protein 1 (PD-1), epidermal growth factor receptor (EGFR) and human epidermal growth factor receptor 2 (HER-2), respectively. Antibody-DNA conjugates (anti-PD1, Cetuximab and Trastuzumab, respectively) were titrated to cells and subsequently labeled with CY5-functionalized imagers (CY5-IM) or CY5-functionalized nanorods (CY5-NR). The mean fluorescent intensities were fitted to a non-cooperative Hill equation and normalized to the fitted maximum fluorescence intensity of cells labeled with a CY5-functionalized imager to extract the apparent dissociation constant. **(c)** Flow cytometric analysis of aPD1-DNA labeled CHO-K1_PD1-high_ cells that were incubated once (single staining) or twice (double staining) with only CY5-IM (blue), only CY5-NR (red) or a combination of both (green). **(d)** Receptor accessibility of CY5-NR as a function of tunable design parameters, including linker length (red), negative charge (green) and DNA handle location (blue). Self-assembly of all DNA nanostructures, including the K_10_-PEG_5K_ and spermine coating, was confirmed using electrophoretic mobility analysis (**Supplementary Figs. 12, 14** and **15**). Individual data points represent the normalized mean fluorescent intensity of 2000 gated single-cell events (n = 3 technical replicates).

To exclude the possibility that the decrease in fluorescent intensity was a result of DNA-antibody or receptor dissociation, ssDNA imager internalization or DNA nanorod impurities (e.g. anti-handle excess still present after NR purification) we performed two additional control experiments. First, unlabeled CHO-K1_PD1-high_ cells that were incubated with CY5-IM or CY5-NR showed similar levels of fluorescent intensity, confirming that CY5-IM and CY5-NR internalization did not impact observed differences in fluorescent intensity (**Fig. 2c**, grey circles). Subsequently, we simultaneously assessed DNA-antibody/receptor dissociation and the influence of DNA nanorod impurities by introducing an additional washing and labeling step. In this experiment we incubated DNA-aPD1 labeled CHO-K1_PD1-_ high cells once or twice with CY5-IM or CY5-NR. Measuring the fluorescent intensity after 1-step labeling or 2-step labeling did not reveal a decrease in fluorescent intensity for CHO-K1_PD1-high_ cells labeled only with CY5-IM or CY5-NR, indicating no apparent dissociation of the antibody (**Fig. 2c**, blue and red circles). Simultaneously, incubating DNA-aPD1 labeled CHO-K1_PD1-high_ cells first with CY5-NR followed by CY5-IM displayed a fluorescent intensity similar to cells incubated with only CY5-IM (**Fig. 2c**, compare blue and green circles), confirming that CY5-NR binds only a fraction of available receptor binding sites. Collectively, these results illustrate that the absolute number of cellular surface receptors targeted by DNA nanorods is limited and comprises only a small fraction of all target receptors present on the cellular surface.

### Determinants of cellular binding efficiency

The limited cellular binding efficiency of DNA nanorods encouraged us to explore determinants of NR that play an important role in receptor binding. First, we focused on the high local concentration of negative charges in the DNA nanostructure caused by phosphate groups in the DNA backbone that might induce electrostatic repulsion in proximity to the negatively charged cell surface. To counteract the overall negative charge, we coated NRs with a polyethylene glycol-oligolysine co-polymer which contains 10 repeats of lysines conjugated to a 5 kDa PEG molecule (K_10_-PEG_5K_). This method has been previously employed to prevent degradation of DNA nanostructures in low-salt conditions and protection from nucleases.^36^ Coating NRs with K_10_-PEG_5K_, however, resulted in decreased cell surface accessibility compared to uncoated NRs (**Fig. 2d**, green circles and **Supplementary Fig. 12**). We hypothesized that this was a result of decreased ssDNA handle availability due to the relatively large PEG molecules on the NR surface. This was supported by additional control experiments using K_10_-PEG_5K_ coated aPD1-NRs which suffered less from decreased binding efficiency compared to uncoated structures (**Supplementary Fig. 13**). Alternatively, we used spermine for NR coating, however this also did not result in improved NR receptor binding (**Fig. 2d**, green circles and **Supplementary Fig. 12**). Taken together, these experiments revealed that counteracting the overall negative charge did not enhance binding efficiency of DNA nanorods, excluding electrostatic repulsion as a major determinant in cellular binding.

Taking advantage of the inherent programmability of DNA origami we next assessed the influence of DNA handle length and handle location. To this end, multiple NRs were self-assembled using DNA handles that contain a 0, 8, 16 and 32-nucleotide (nt) single-stranded linker that separates the antibody from the NR surface (**Supplementary Fig. 14** and **Supplementary Table. 4**). Unsurprisingly, NRs that contained DNA handles with a 32-nt spacer showed the highest binding efficiency, however, the increase in cellular binding was only moderate compared to other linker lengths (**Fig. 2d**, red circles). In addition, fortification of the 16-nt and 32-nt linker using a complementary anti-handle did not improve binding efficiency compared to single-stranded linkers. Finally, we constructed four unique NR configurations in which the position of the ssDNA handles was varied (**Fig. 2d**, right bottom, **Supplementary Table. 4**). Since DNA handle incorporation efficiency strongly correlates with the position in the structure^48^, DNA handle incorporation for each configuration was quantified. Using gel mobility electrophoresis, we demonstrated that configuration *1*, *3* and *4* had similar handle incorporation efficiency while configuration 2 suffered from missing DNA handles (**Supplementary Fig. 15**). Despite the lower incorporation efficiency, configuration *2* as well as configuration *3* showed improved cellular binding efficiency compared to configuration *1* and *4* (**Fig. 2d**, blue circles). We attribute this improved binding efficiency to the relative orientation of NRs with respect to the cell membrane and therefore the accessibility of tethered antibodies to the cell-bound receptors. More specifically, configuration *1* and *4* would result in lateral receptor engagement whereas configurations *2* and *3* facilitate axial receptor binding. These results demonstrate that receptor binding efficiency can be modulated using DNA handle location and strongly correlates with NR orientation during receptor engagement.

### DNA nanostructure size and shape influence cellular binding efficiency

Encouraged by the observed relation between DNA nanorod orientation and cellular binding efficiency, we constructed multiple DNA nanostructures to evaluate the effect of shape and size on receptor binding. Two additional nanostructures, a twist-corrected rectangular DNA origami rectangle^20,49^ (Rec, 75×100 nm^2^) and a tetrahedral DNA nanostructure^50^ (Tet), respectively, were successfully folded and purified (**Fig. 3a** and **Supplementary Fig. 16** and **17**). Additionally, a 50-nt double-stranded binding probe (dsP) was self-assembled. The DNA nanorod and DNA rectangle contained two ssDNA handles which facilitate binding to a single antibody (**Supplementary Fig. 18**). In contrast, the DNA tetrahedron and double-stranded probe employed a single ssDNA handle which allows binding of two nanostructures to a single antibody, comparable to an imager strand (**Fig. 2a**). Consequently, double-stranded binding probes and tetrahedrons are therefore labeled with a single CY5 fluorophore, while DNA rectangles and nanorods contain two CY5 labels. To accurately compare binding efficiency, CY5 fluorescent intensity of all nanostructures should be similar and scale proportionally with the number of dyes incorporated (e.g. the fluorescent intensity of the double-stranded binding probe or tetrahedron should be two times smaller than the fluorescent signal of DNA rectangles or nanorods). Using fluorescent intensity measurements and a fixed concentration of CY5-functionalized nanostructures, we demonstrated similar fluorescent intensities for CY5-IM, CY5-dsP and CY5-Tet as well as for CY5-Rec and CY5-NR, however, it also revealed a non-proportional increase (~3-fold) in fluorescent intensity when comparing CY5-IM, CY5-dsP or CY5-Tet to CY5-Rec and CY5-NR (**Supplementary Fig. 19**). As a result, receptor binding efficiencies of the DNA rectangles or nanorods could be slightly over-estimated when compared to the single stranded imager, double-stranded probe or tetrahedron. However, since the observed difference in fluorescent intensity between the large DNA nanostructures (i.e. CY5-Rec or CY5-NR) and the small DNA probe is >10-fold, we decided not to use a correction factor for this over-estimation and directly rely on the fluorescent intensity observed using flow cytometric analysis. Incubating all DNA nanostructures with aPD1-DNA labeled CHO-K1_PD1-high_ cells revealed a negative correlation between nanostructure size and receptor binding efficiency. The binding efficiency of DNA rectangles, comprising a larger surface area than DNA nanorods, showed the lowest binding efficiency indicating that aspect ratio and surface area are critical determinants of DNA nanostructure binding (**Fig. 3b**, compare blue circles). Recovery of the fluorescent intensity after incubating labeled cells with CY5-imager strands confirmed that the decreased fluorescent intensity was a result of limited DNA nanostructure binding, rather than DNA-antibody dissociation or DNA handle impurities (**Fig. 3b**, red circles). These results collectively demonstrate that nanostructure size negatively impacts cellular binding and suggest aspect ratio and surface area as potential parameters that modulate receptor binding efficiency.

**Figure 3.**
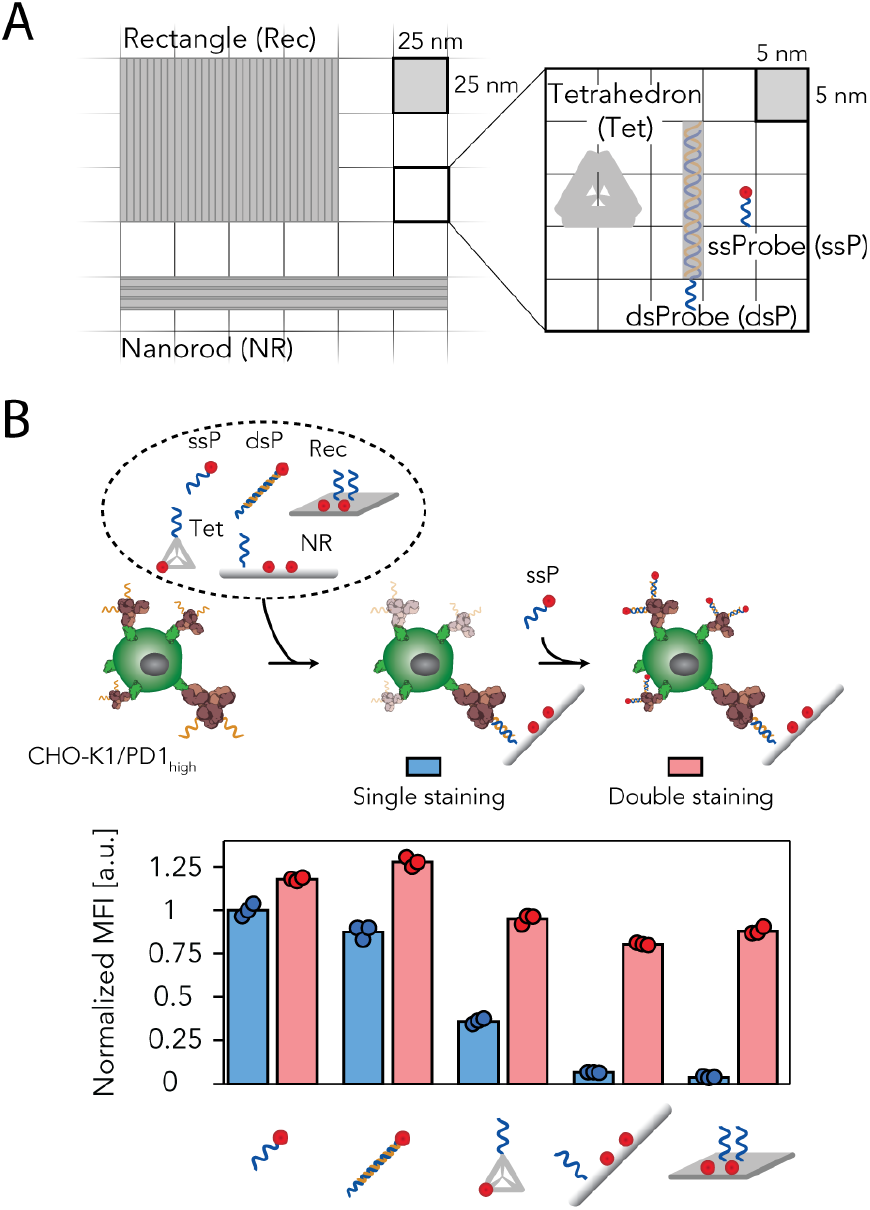
Effect of DNA nanostructure size and shape on receptor accessibility. **(a)** Schematic overview of the dimensions of different DNA nanostructures and smaller DNA probes used to target cellular surface receptors (see also **Supplementary Fig. 16** and **17**). **(b)** Flow cytometric analysis of aPD1-DNA labeled CHO-K1_PD1-high_ cells first incubated with 20 nM CY5-functionalized DNA nanostructures or probes for 30 min (Single staining, blue) followed by incubation with 20 nM CY5-functionalized imagers for 30 min (Double staining, red). Individual data points represent the normalized mean fluorescent intensity of 2000 gated single-cell events (n = 3 technical replicates).

### Cellular determinants of DNA nanostructure binding efficiency

We have demonstrated that DNA nanostructure size limits cell surface accessibility and that handle location is a critical parameter to modulate receptor binding; however, all these determinants are intrinsically related to the DNA nanostructure. We therefore next investigated how cellular features, such as surface receptor density or the presence of a dense glycocalyx, impact receptor targeting efficiency of DNA nanostructures. First, we assessed the effect of PD1 density on nanostructure binding. We hypothesized that over-expression of target receptors could result in a receptor density that exceeds the theoretical number of DNA nanostructures that can bind based on surface area. Consequently, receptor binding of an individual nanostructure could block access to other surface receptors limiting overall availability of receptor binding sites. To study this effect, we analyzed binding of CY5-IM and CY5-NR to CHO-K1 cells expressing low, intermediate, and high levels of PD1 (**Fig. 4a** and **Supplementary Fig. 10**). Unsurprisingly, CHO-K1 cells incubated with CY5-IM displayed a decrease in fluorescent intensity as a function of receptor density (**Fig. 4a**, compare blue circles). Moreover, the fluorescent intensity of CHO-K1_PD1-low_ cells labeled with CY5-IM exceeds that of CHO-K1_PD1-high_ cells labeled with CY5-NR. This indicates that even CHO-K1_PD1-low_ cells express sufficient PD1 receptors to reach similar fluorescent intensity levels as CHO-K1_PD1-high_ cells when incubated with CY5-NR. Specifically, if over-expression of target receptors significantly interferes with DNA nanostructure binding, the fluorescent intensity of CHO-K1_PD1-low_ labeled with CY5-NR should be comparable to that of CY5-NR labeled CHO-K1_PD1-high_ cells. Analyzing binding of CY5-NR to CHO-K1 cells, however, displayed a decreasing trend in fluorescent intensity as a function of PD1 expression comparable to CY5-IM, providing direct evidence that PD1 receptor density does not play a major role in DNA nanostructure binding (**Fig. 4a**, compare red circles). In addition to target receptor density, we also explored whether the presence of other surface proteins present in the crowded environment of the cell membrane could interfere with DNA nanostructure binding. To examine this, we analyzed binding of CY5-IM and CY5-NR to CHO-K1_PD1-high_ cells that were dissociated using the proteolytic enzyme trypsin. Treating cells with trypsin resulted in a 6.8-fold difference in fluorescent intensity between CY5-IM and CY5-NR compared to a 17.2-fold difference observed for untreated cells, indicating that the crowded cellular surface limits DNA nanostructure binding (**Fig. 4b**). Taken together, these results illustrate that the presence of other cellular surface proteins negatively impacts receptor accessibility while the target receptor density has only a minor contribution.

**Figure 4.**
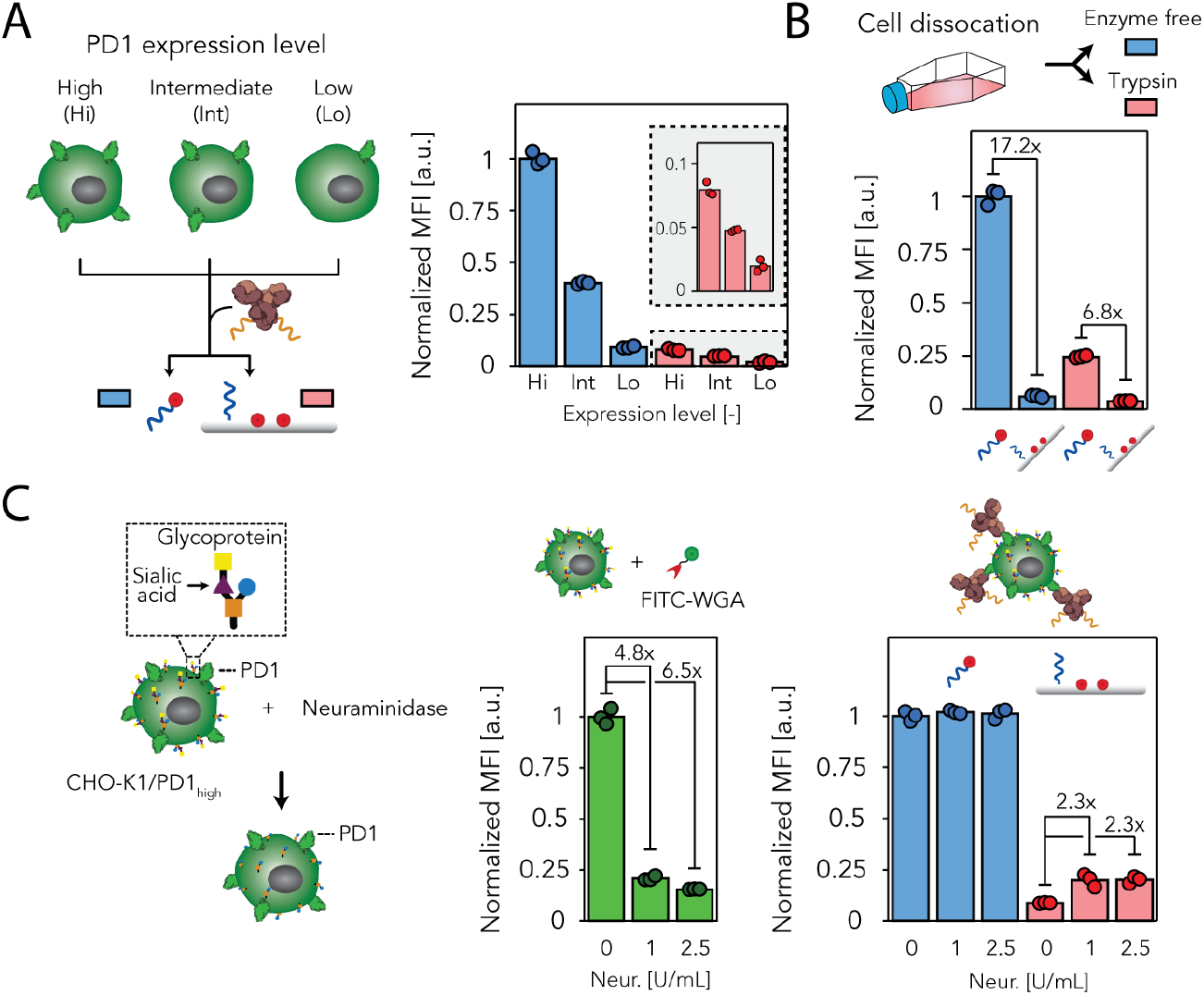
Effect of cellular determinants on DNA nanostructure-receptor interaction. **(a)** Flow cytometric analysis of CHO-K1_PD1-high_ cells expressing different levels of PD1 (High, Hi; Intermediate, Int; Low, Lo). Cells were labeled with aPD1-DNA followed by incubation with CY5-functionalized imager or CY5-functionalized DNA nanorod. **(b)** Flow cytometric analysis of CHO-K1_PD1-high_ cells that were dissociated using an enzyme-free dissociation buffer or trypsinization or **(c)** treated with different concentrations of neuraminidase (Neur.) to remove sialic acids. Cellular labeling was performed as described in **(a).** Fluorescein isothiocyanate-linked wheat germ agglutinin (FITC-WGA) was used to detect sialic acid residues. Individual data points represent the normalized mean fluorescent intensity of 2000 gated single-cell events (n = 3 technical replicates).

Having established that receptor accessibility is sensitive to the presence of cellular surface proteins, we sought to examine the impact of the glycocalyx on DNA nanostructure binding. Previous research has shown that enzymatic digestion of the glycocalyx resulted in enhanced nanoparticle uptake.^51,52^ To assess the effect of the glycocalyx on DNA nanostructure receptor binding, CHO-K1_PD1-_ high cells were treated with neuraminidase to selectively remove sialic acids (**Fig. 4c**). CHO-K1_PD1-high_ cells treated with neuraminidase and incubated with CY5-IM did not exhibit improved labeling efficiency, indicating that the small imager is not affected by glycocalyx composition. In contrast, cells incubated with CY5-NR displayed a 2.3-fold increase in fluorescent intensity, confirming that the glycocalyx interferes with DNA nanostructure binding. Overall, these findings indicate that receptor targeting efficiency of DNA nanostructures is not only dependent on nanostructure design but is also significantly impacted by crowding on the cell membrane.

### aPD1-functionalized DNA nanorods do not block immune checkpoint receptors

Finally, to correlate cellular binding efficiency and modulation resultant downstream signaling, we explored how limited cellular binding efficiency of DNA nanostructures translates to receptor blocking efficiency *in vitro*. As receptor blocking efficiency is pivotal to effective immunotherapy^44^, we hypothesized that low receptor binding efficiency of aPD1-functionalized nanorods could result in decreased checkpoint blockade. For these studies, we used a commercially available bioassay to measure the ability of aPD1 to block PD1/PDL1 interactions based on Jurkat T_PD1/TCR_ cells reporters and CHO-K1_PDL1/APC_ cells as antigen presenting cells (**Fig. 5a**). When aPD1 antibodies were titrated to a coculture of Jurkat T_PD1/TCR_ cells and CHO-K1_PDL1/APC_, Jurkat T_PD1/TCR_ cells responded in a dose-dependent manner, indicating inhibitory activity of PD1 signaling in this cell system (**Supplementary Fig. 20**). Before assessing the blocking efficiency of aPD1-functionalized nanorods we performed additional control experiments to validate the purity and structural integrity of DNA nanorods in culture medium. After aPD1-functionalization of DNA nanorods we employed 2 rounds of PEG precipitation^43,53^ or agarose gel extraction^46^ to remove free aPD1 and found that only agarose gel purification resulted in full removal of free aPD1 antibodies (**Supplementary Fig. 21** and **22**). Additionally, we confirmed the stability of aPD1-functionalized nanorods in culture medium for the duration of the blocking assay (**Supplementary Fig. 23**). Treating a coculture of Jurkat T_PD1/TCR_ cells and CHO-K1_PDL1/APC_ with IgG isotype control, empty-NR, aPD1-NR and free aPD1 showed that only free aPD1 was able to block PD1/PDL1 signaling (**Fig. 5b**). Moreover, no significant difference in receptor blocking efficiency was found between empty-NR and aPD1-NR, which is in line with recent work of Fang *et al.*^28^ that showed that a single PDL1 ligand presented on a DNA origami flatsheet did not elicit PD1 inhibition. These results illustrate that inefficient cellular targeting of antibody functionalized DNA nanostructures directly translates to decreased receptor blocking efficiency *in vitro*.

**Figure 5.**
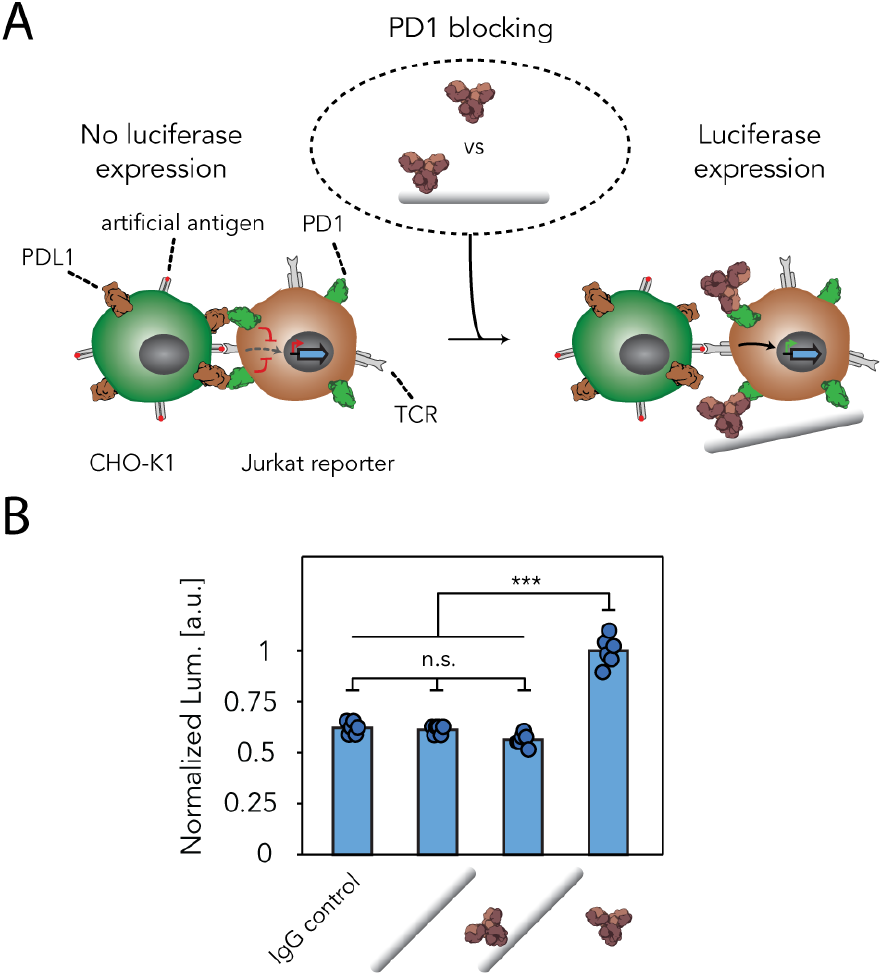
aPD1-functionalized DNA nanostructures are inefficient immune checkpoint inhibitors. **(a)** Schematic of the cell assay to detect the potency of aPD1-functionalized DNA nanorods (aPD1-NR) to block PD1/PDL1 interactions. Specifically, artificial antigen-presenting (aAPC) CHO-K1_PDL1/APC_ cells that express the programmed death-ligand 1 (PDL1) were cocultured with Jurkat T_PD1/TCR_ cells stably expressing PD1, T-cell receptors (TCRs) and a luciferase induced by nuclear factor of activated T cells (NFAT). **(b)** Luciferase expression from a CHO-K1_PDL1/APC_ /Jurkat T_PD1/TCR_ cell coculture treated with IgG, empty-NR, aPD1-NR or free aPD1. One-way analysis of variance (ANOVA) was used followed by Tukey’s multiple-comparison test (\*\*\**P* < 0.001). Individual data points represent normalized luminescence (Lum.). Experiment was performed independently in duplicate with 3 technical replicates for each experiment.

## Discussion

DNA nanotechnology has facilitated the design of a library of nanostructures that have been shown to be stable in cellular environments and can be readily modified with small molecules or protein ligands to study cellular signaling at the nanoscale or act as programmable delivery vehicles.^54,55^ Here, we evaluated receptor binding efficiency of antibody-functionalized DNA nanostructures to elucidate critical design parameters that can promote or hamper cellular binding. Our results reveal that, while the native affinity of incorporated antibodies remains unaltered, the absolute number of surface receptors targeted by antibody-functionalized DNA nanostructures is reduced compared to free antibodies. Systematic evaluation of nanostructure design parameters revealed that nanostructure orientation and size are key parameters for efficient receptor binding and demonstrated that the cell surface composition acts as a natural barrier that limits receptor accessibility. Based on these findings we hypothesize that steric hindrance caused by the larger DNA nanostructures is the primary contributor to limited receptor binding efficiency. A potential application of this nanostructure induced steric hindrance could comprise the formation of a steric barrier around the cell that is able to block all ligand-receptor interactions. However, experimental evidence showed that binding of large DNA nanostructures to surface receptors did not prevent other, smaller, probes from accessing unbound receptors, excluding the possibility of using large DNA nanostructures as tools for cell signaling blockage. Moreover, a cellular assay that assessed immune checkpoint blockade displayed that decreased receptor binding efficiency of aPD1-nanostructures directly translated to limited blocking of immune checkpoint receptors. This result highlights important considerations for the use of nanostructures in biological systems and their therapeutic effectiveness. For example, smaller nanostructures containing only a limited number of therapeutic antibodies might be beneficial over larger nanostructures that contain multiple antibodies to block ligand-receptor signaling. Rationalizing the impact of individual design parameters for ligand-functionalized DNA nanostructures therefore provides a powerful addition to the design criteria for nanostructures targeting cellular surface receptors.

Aside from the absolute number of receptors that are targeted, cellular activation mechanisms also play a major role in cellular signaling. Previous work, which employed DNA nanostructures to study distance dependent effects of receptor activation with nanometer precision, showed that ligand-functionalized DNA origami structures induced similar or even enhanced cellular signaling compared to soluble ligands.^24,26–28^ Combining these results with the findings in our work suggests that cellular signaling mostly depends on the activation mechanism, rather than the number of receptors that are bound. This underlines the limitations of using a single parameter, the equilibrium dissociation constant (K_D_), to assess ligand-functionalized nanostructure performance.^56^ Dissociation constants only refer to the strength of individual ligand-receptor interactions excluding the effect of nanostructure design or cellular determinants. Our work shows that the efficacy of cellular targeting is dictated by a combination of receptor affinity and accessibility of receptors at the target site. As such, a broader subset of parameters, which include cell signaling modulation and receptor binding efficiency, should be explored to maximize the potential of nanomedicines. We envision that programmable DNA nanostructures find great application in the elucidation on critical design parameters that will eventually guide the design of precision medicines, either composed of nucleic acids, polymers or organic molecules.

## Author Contributions

G.A.O.C. designed the study, performed experiments, analyzed the data, and wrote the manuscript. B.J.H.M.R. performed and analyzed the AFM imaging, analyzed the data, and wrote the manuscript. A.M. performed and analyzed the TEM imaging. N.B.T developed and derived the thermodynamic model. S.M.J.v.D and H.v.E. developed and provided the stable CHO-K1 cell lines and provided critical input for the experiments. L.A. analyzed the data and wrote the manuscript. T.F.A.d.G. conceived, designed and supervised the study, analyzed the data and wrote the manuscript. All authors gave approval to the final version of the manuscript.

## Notes

The authors declare no competing financial interest.

## Acknowledgements

We thank J. Schill for help with the TEM imaging and analysis. M. Merkx and P. de Vink are gratefully acknowledged for valuable insights and fruitful discussions. We thank Promega for making it possible to provide the PD1/PDL1 Blockade Bioassay cells. This work was supported by the European Research Council (ERC) (project no. 677313 BioCircuit), an NWO-VIDI grant from The Netherlands Organization for Scientific Research (NWO, 723.016.003), and funding from the Ministry of Education, Culture and Science (Gravity programs, 024.001.035 & 024.003.013)

