## Supplementary Information for "Determinants of ligand-functionalized DNA nanostructure-cell interactions"

### Supplementary methods

##### DNA and protein sequence protein G

The sequence of protein G as previously described in Cremers *et al.* [1]. The single-letter amino acid code is shown in uppercase above the corresponding DNA sequence. The cysteine is shown in purple, the *strep*-tag in orange, protein G (pG) in blue, the hexahistidine tag in green and the amber stop codon, coding for the non-natural amino acid *p*-Bpa, is indicated in red.

```
        M  S  C  W  S  H  P  Q  F  E  K  G  T  M  T  F  K  L  I  I
1       ATGAGTTGCTGGTCCCATCCGCAGTTCGAGAAAGGTACCATGACATTTAAACTGATAATC

        N  G  K  T  L  K  G  E  I  T  I  E  A  V  D  A  *  E  A  E
61      AACGGCAAAACCTTAAAAGGGGAGATCACAATTGAGGCAGTCGATGCCTAGGAAGCCGAG

        K  I  F  K  Q  Y  A  N  D  Y  G  I  D  G  E  W  T  Y  D  D
121     AAAATCTTTAAACAATATGCTAATGATTATGGTATTGACGGAGAATGGACGTATGACGAT

        A  T  K  T  F  T  V  T  E  E  F  T  S  G  G  S  G  D  D  H
181     GCGACAAAAACTTTCACCGTAACTGAGGAATTCACTAGTGGTGGAAGTGGGGACGATCAT

        H  H  H  H  H  *
241     CATCATCATCATCATTAA
```

##### Recombinant protein expression and purification

To express protein G we used the exact same method as previously described in Cremers *et al.* [1]. A pET28a(+) vector encoding the pG gene as reported previously[1] was co-transformed with the pEVOL-pBpF vector, encoding for the orthogonal aminoacyl tRNA synthetase/tRNA pair (kindly provided by Peter Schultz), into *E. coli* BL21(DE3) (Novagen) for protein expression. A single colony of freshly transformed cells was cultured at 37 °C in 500 mL 2xYT medium supplemented with 50 µg/mL kanamycin (Merck) and 25 µg/mL chloramphenicol (Sigma Aldrich). When the $OD_{600}$ of the culture reached ~0.6, protein expression was induced by addition of β-D-1-thiogalactopyranoside (IPTG, Applichem), arabinose (Sigma Aldrich) and the unnatural amino acid *p*-Benzoyl-L-phenylalanine (*p*-BpA) (Bachem) in a final concentration of 1 mM, 0.02% (w/v) and 1 mM, respectively. The induced protein expression was carried out for ~18h at 25 °C and subsequently the cells were harvested by centrifugation at 10,000 xg for 10 min at 4 °C. The cell pellet was resuspended in BugBuster (5 mL/g pellet, Merck) supplemented with benzonase 5 (µL/g pellet, Merck) and incubated for 45 minutes on a shaking table. The suspension was centrifugated at 40,000 xg for 30 min at 4 °C and the supernatant was subjected to Ni-NTA affinity chromatography on a gravity column. His-tagged pG was loaded on the column and washed with washing buffer (1x PBS, 370 mM NaCl, 10% (v/v) glycerol, 20 mM imidazole, pH 7.4) before elution with his-elution buffer (1x PBS, 370 mM NaCl, 10% (v/v) glycerol, 250 mM imidazole, pH 7.4). Subsequently, the Ni-NTA elution fraction was loaded on a *Strep*-tactin column, washed with washing buffer (100 mM Tris-HCl, 150 mM NaCl, 1 mM EDTA, pH 8.0) and eluted using wash buffer supplemented with 2.5 mM desthiobiotin (IBA Life Sciences). Proteins were stored at -80 °C in 1 mL aliquots of 50 µM at -80°C in a buffer containing 100 mM Tris-HCl, 150 mM NaCl, 1 mM EDTA and 2 mM TCEP at pH 8.0. The concentration of pG was calculated based on the absorption at 280 nm (ND-1000, Thermo Scientific) assuming an extinction coefficient of 15,470 $M^{-1}$ $cm^{-1}$ and the purity of pG was assessed on reducing SDS-PAGE.

**Purification of antibody-DNA conjugates using HPLC**

Size exclusion chromatography was performed on an Agilent 1260 Infinity II Bio-inert LC System using a Bio-SEC-5 column (5 µm, 300 Å, 7.8*300 mm, Agilent). The column was equilibrated with 5 column volumes of 1x PBS, pH 7.2 using a flow rate of 1 mL/min. The column was coupled to a 280 nm UV spectrophotometer and a fraction collector. 95 µL sample was injected and elution fractions of 0.2 mL were collected and analyzed on SDS-PAGE under reducing conditions

**Gel electrophoresis**

For SDS-PAGE, samples were heated to 95°C for 5 min in loading buffer (20 mM Tris, 3.33% glycerol, 0.83% SDS (w/v) , 0.003% bromophenol blue, pH 6.8) and 50 mM of DTT was added when the gel was run under reducing conditions. SDS-Page was performed using pre-cast 4-20% Mini-PROTEAN® TGX gels (Bio-rad) in running buffer (25 mM Tris, 192 mM glycine, 0.1% SDS, pH 8.3) for 45-50 min at 150 V. As protein reference the Precision Plus Protein All Blue ladder (Bio-Rad) was used. Eventually, gels were stained with Coomassie Brilliant Blue G-250 (Bio-Rad), imaged using an ImageQuant 400 Digital Imager (GE Healthcare), and analyzed with ImageJ.

Native-PAGE analysis was used to confirm self-assembly of the tetrahedron and dsProbe (**Supplementary Figure 13**). Gels were prepared at 6% monomer concentration (30%, 19:1 stock concentration acrylamide) in gel buffer (0.5x TBE, 10 mM $MgCl_2$, pH 8.0). Samples were diluted in sample buffer to a final concentration of (10% Sucrose, 0.5x TBE, 10 mM $MgCl_2$, pH 8.0). The gel was run in 0.5x TBE, 5 mM $MgCl_2$, pH 8.0 for 20 min at 200V in an ice bath and post-stained with SYBR Gold (Thermo Scientific). GeneRuler Ultra Low Range DNA ladder (Thermo Scientific) was included as a reference ladder.

Agarose gel electrophoresis was used to confirm DNA origami self-assembly. Briefly, 1.5% agarose gels were cast in gel buffer (0.5x TBE, 10 mM $MgCl_2$, pH 8.0) supplemented with SYBR Safe. Gels were run in gel buffer for 90 min at 65 V in an ice bath. DNA origami samples were loaded using Ficoll-400 (final concentration 1.5% (w/v)). Gels were imaged using an ImageQuant 400 Digital Imager (GE Healthcare) and analyzed with ImageJ.

**PEG precipitation to remove excess staple strands**

Excess staple strands were removed by 2 rounds of PEG precipitation.[2] Briefly, a precipitation buffer containing a final concentration of 15% PEG8000 (w/v), 5 mM Tris-HCl, 1 mM EDTA and 505 mM NaCl (pH 8.0) was prepared and mixed in a 1:1 ratio with the DNA origami reaction mixture. The solution was mixed and spun at 21,000 xg, at 16 °C for 25 min using a microcentrifuge. The supernatant was removed and the pellet was redissolved in folding buffer (5 mM Tris, 1 mM EDTA, 25 mM NaCl, 12 mM $MgCl_2$) over the course of 30 min at 30 °C. The cycle was repeated once more and eventually the pellet was dissolved in 1x PBS, pH 7.4 supplemented with 10 mM $MgCl_2$.

**PEG precipitation to remove free aPD1-DNA conjugates**

A precipitation buffer containing a final concentration of 15% PEG8000 (w/v), 5 mM Tris-HCl, 1 mM EDTA and 418 mM NaCl (pH 8.0) was prepared and mixed in a 1:1 ratio with the DNA origami reaction mixture. The solution was mixed, incubated for 10 min at 4°C and spinned at 16,000 xg, at 4 °C for 25 min using a microcentrifuge. The supernatant was removed and the pellet was redissolved in 1x PBS, pH 7.4 supplemented with 10 mM $MgCl_2$ over the course of 30 min at 25 °C. The cycle was repeated once more.

**Agarose gel extraction**

Agarose gel extraction was performed as described previously.[3] Specifically, an agarose gel casting tray (7x10 cm) was filled with 20 mL of 4% agarose in gel buffer (0.5x TBE, 10 mM $MgCl_2$, pH 8.0). After solidifying, a second layer was added using 45 mL 1.5% agarose in gel buffer supplemented with SYBR Safe. DNA origami samples were loaded using Ficoll-400 (final concentration 1.5% (w/v) and run for 90 minutes at 65V in an ice bath. After running, a band just below the DNA origami was excised and the band was replaced with an extraction buffer (30% sucrose (w/v), 0.5x TBE, 10 mM $MgCl_2$, pH 8.0). The gel was run for another 30 minutes to electrophorese the DNA nanostructure in the extraction buffer. The extraction buffer was recovered, and the purified DNA origami solution was concentrated using PEG precipitation with a precipitation buffer containing 20% PEG8000 (w/v), 5 mM Tris-HCl, 1 mM EDTA and 505 mM NaCl (pH 8.0).

**AFM imaging of DNA nanostructures**

Topographic images were acquired in tapping mode under liquid conditions on a MultiMode 8 atomic force microscope with a NanoScope IIIa controller (Veeco) using V-shaped $Si_3N_4$ cantilevers with sharpened pyramidal tip and a nominal spring constant of 0.04 N/m (OTR4, Bruker AFM Probes). Substrates were prepared by attaching laser-cut mica discs (~1 $cm^2$ , Ted Pella) to Teflon (VWR) using epoxy glue. DNA origami solutions were diluted to 2 nM with imaging buffer (10 mM Tris, 1 mM EDTA, 10 mM $MgCl^2$, pH 8.0) and 5 µL was deposited on a freshly-cleaved mica substrate. The sample was incubated for 30 s and subsequently 50 µL of imaging buffer was added. Images (512×512 px) of 1.2×1.2 $\mu m^2$ or 2.0×2.0 $\mu m^2$ were acquired, optimizing the scanning and feedback parameters for each image. All images were analyzed using Gwyddion v2.39 software.

### Supplementary figures

#### Expression of protein G (pG)

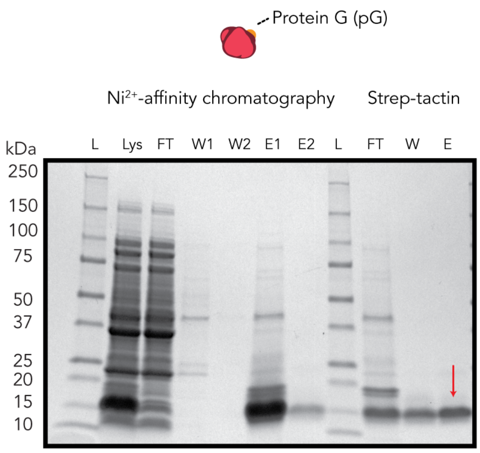

**Supplementary Figure 1 | SDS-PAGE analysis of the expression and purification of protein G**. Image of a SDS-PAGE gels (4-20% polyacrylamide, stained for proteins with Coomassie Brilliant Blue), confirming successful expressing and purification of protein G (pG, calculated mass: 9784.64 Da). pG runs at an apparent mass of 14 kDa. Labels: L, reference protein ladder; Lys, bacterial lysate; FT, flow through; W, wash; E, elution.

#### Purification and incorporation of aPD1-DNA conjugates

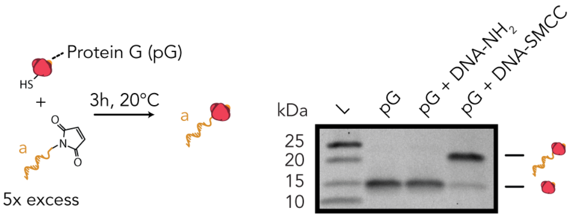

**Supplementary Figure 2 | SDS-PAGE analysis of a protein G-DNA conjugation reaction**. Image of a SDS-PAGE gels (4-20% polyacrylamide, stained for proteins with Coomassie Brilliant Blue), of the conjugation process for protein G (pG) to sulfosuccinimidyl 4-(N-maleimidomethyl)cyclohexane-1-carboxylate (Sulfo-SMCC) functionalized p1 (5 molar excess, ODN mass, 6.4 kDa). DNA conjugation to the thiol functionality located N-terminally in pG results in a selective gel shift of the band corresponding to pG. Label: L, reference protein ladder.

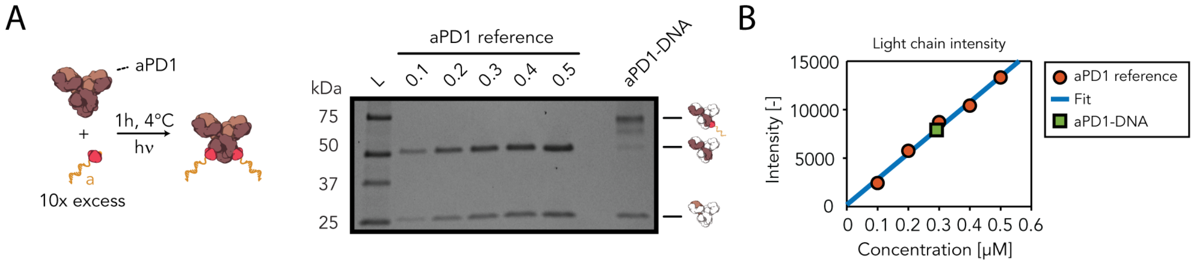

**Supplementary Figure 3 | SDS-PAGE analysis of an antibody protein G-DNA conjugation and purification reaction**. **(a)** Image of a reduced SDS-PAGE gel (4-20% polyacrylamide, stained for proteins with Coomassie Brilliant Blue) of the conjugation process for protein G-DNA to an anti-PD1 antibody. Protein G-DNA (10 molar equivalents) conjugation to the Fc region of the antibody for 1h at 4 °C using 365 nm illumination results in a gel shift of the band corresponding to the heavy chain of the antibody. Since pG-DNA conjugates were not purified before coupling, a second band just below the final product is apparent which represents a heavy chain coupled to pG without a DNA handle. A calibration dilution series (0.1-0.5 µM) of aPD1 is used to accurately estimate the concentration of conjugated DNA-aPD1. **(b)** Gel densitometry of the antibody light chain is used to accurately determine the concentration of DNA-aPD1 conjugates by comparing the gel band intensity to a calibration dilution series (0.1-0.5 µM) of aPD1. ImageJ was used to determine the light chain band intensity (area under the curve). The red circles indicate the optical density of the calibration dilution series and are fitted to a linear regression model. The calibration curve was used to determine the concentration of DNA conjugated aPD1. Label: L, reference protein ladder.

#### Incorporation of aPD1 on DNA nanorod

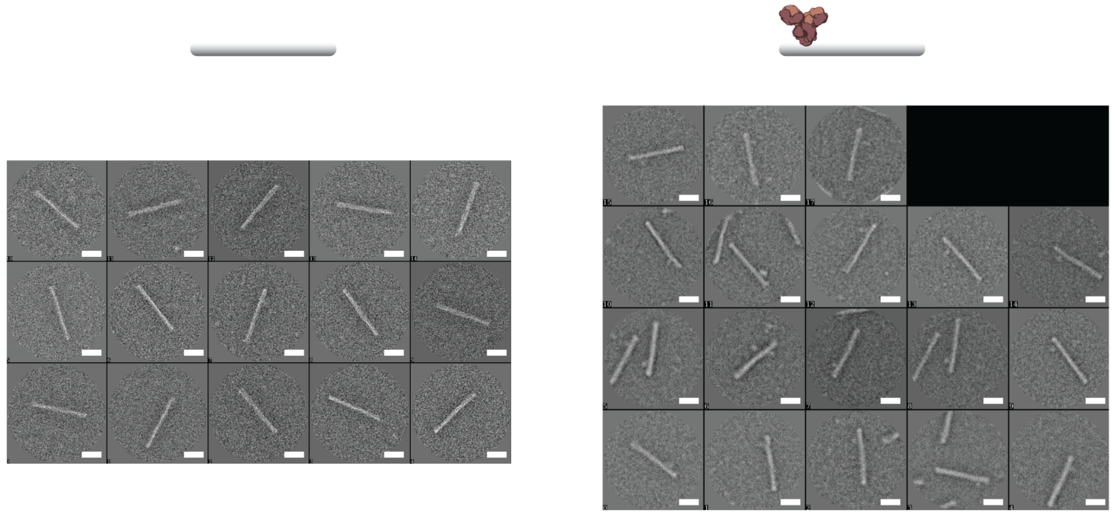

**Supplementary Figure 4 | TEM analysis.** Single-particle TEM micrographs of DNA origami nanorods without aPD1 functionalization (left) or with aPD1 (right) used for reference-free class averages as depicted in **Figure 1b**. 2.5 nM nanorods were stained with 0.4% (w/v) aqueous uranyl acetate solution onto a carbon coated grid (Cressington 206Carbon). Scale bar, 50 nm.

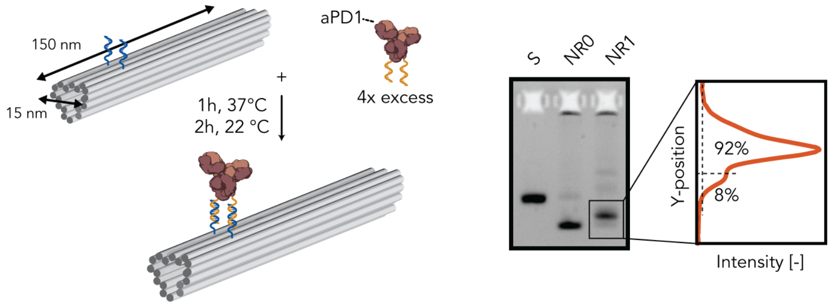

**Supplementary Figure 5 | Structural integrity of DNA nanorods functionalized with aPD1**. Image of an agarose gel (1.5% agarose, stained for DNA with SYBR safe) of a self-assembled nanorod with and without DNA-aPD1 conjugate. DNA nanorods functionalized with two DNA handles were incubated with 4 molar equivalents of aPD1-DNA for 1h at 37 °C followed by 2h at 22 °C. aPD1 incorporation efficiency was assessed using gel densitometry. ImageJ was used to quantify the area under the curve to estimate the fraction of nanorods that were functionalized with aPD1 (top band). Using this analysis, we found an aPD1 incorporation efficiency of ~92%.

#### Characterization of aPD1-nanorod stability and cellular uptake

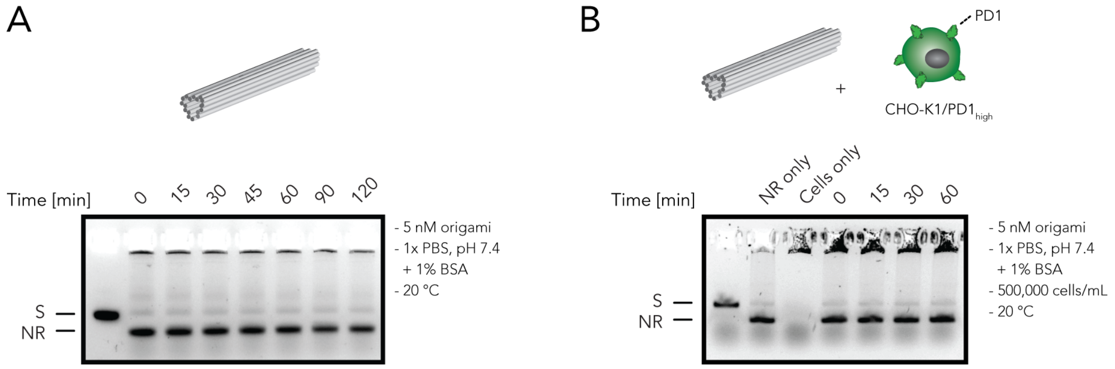

**Supplementary Figure 6 | Structural integrity of DNA nanorods during cellular labeling**. Images of agarose gels (1.5% agarose, stained for DNA with SYBR safe) of a self-assembled nanorod incubated in cellular labeling buffer (1x PBS + 1% BSA, pH 7.4) in the **(a)** absence and **(b)** presence of CHO-K1 cells to monitor stability of DNA nanorods during cellular labeling.

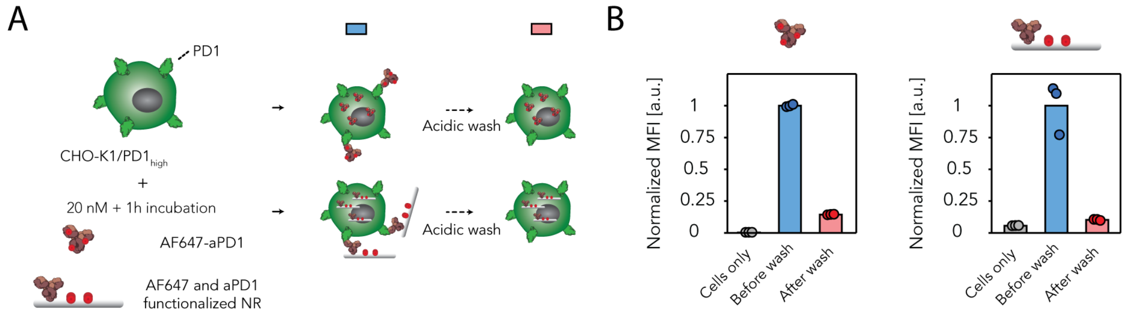

**Supplementary Figure 7 | Internalization of aPD1-NR and free aPD1**. **(a)** Schematic overview of the experimental set-up used to determine the degree of internalization of 20 nM aPD1 after 1h incubation at room temperature. Removal of surface aPD1 with an acidic washing buffer (RPMI1640 + 2% FBS, pH 2.0) for 5 min at 37 °C allows detection of internalized aPD1 using flow cytometry. **(b)** Flow cytometric measurement of CHO-K1$_{PD1\text{-}high}$ cells incubated for 1h with 20 nM of Alexa-647-labeled (AF647) free aPD1 or aPD1-NR before (blue) and after (red) acid wash removal of surface antibody. Acid-stripped CHO-K1$_{PD1\text{-}high}$ cells show similar fractions of internalized AF647-labeled aPD1 or AF647-aPD1-NR, excluding antibody internalization as the primary contributor for the difference in fluorescent intensity levels. The data is normalized to the mean fluorescence intensity of CHO-K1$_{PD1\text{-}high}$ cells labeled with AF647-aPD1 and AF647-aPD1-NR, respectively, measured before the acidic wash. Individual data points represent the normalized mean fluorescent intensity of 2000 gated single-cell events (n = 3 technical replicates).

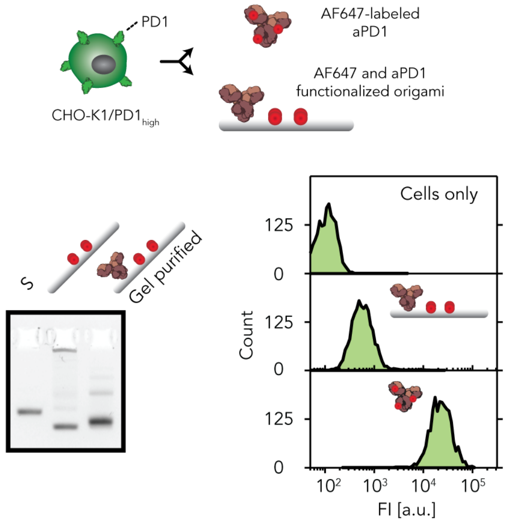

**Supplementary Figure 8 | Cellular binding of agarose gel extracted aPD1-NR nanostructures**. DNA nanorods self-assembled with four Alexa 647 fluorophores and functionalized with a single aPD1 antibody (AF647-aPD1-NR) were purified by loading the reaction mixture on an agarose gel (1.5%), running the gel for 90 min in an ice bath, and extracting the band corresponding to AF647-aPD1-NR as described in the **Methods** section. The image of an agarose gel (1.5% agarose, stained for DNA with SYBR safe) confirms the structural integrity and AF647-aPD1-NR incorporation after gel extraction. CHO-K1$_{PD1\text{-}high}$ cells were incubated with 20 nM AF647-aPD1-NR or free AF647-aPD1 for 60 min at room temperature and fluorescent intensity levels were quantified using flow cytometry.

#### aPD1-AF647 labeling and compensation

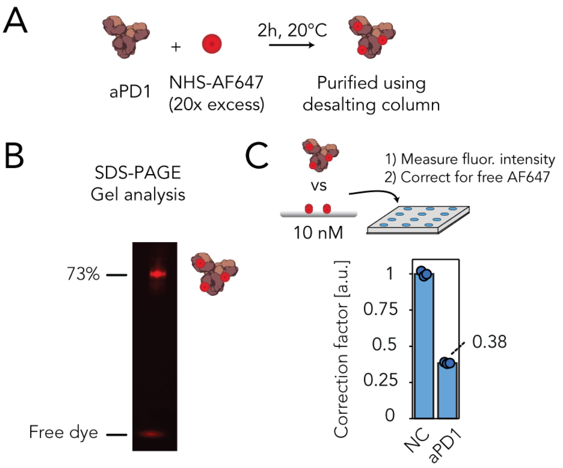

**Supplementary Figure 9 | Compensation for AF647 labeling efficiency of aPD1. (a)** AF647-NHS was coupled to aPD1 by incubating a 20-fold molar excess of AF647-NHS for 2h with aPD1. Non-reacted dye was removed by gel filtration using a ZEBA spin desalting column. **(b)** Image of a non-reduced SDS-PAGE gel (4-20% polyacrylamide, unstained) of the purified AF647-aPD1 antibody. Gel band intensity analysis was used to determine contamination of free AF647. **(c)** Fluorescence intensity measurement of 10 nM AF647-aPD1 and 10 nM of a DNA nanorod functionalized with four AF647 fluorophores. The fluorescent intensity of AF647-aPD1 was corrected for contamination of free AF647 dye (multiplication with 0.73) and the corrected fluorescent intensity was displayed. The correction factor of 0.38 was used to plot **Figures 1d** and **1e**. Individual data points represent the fluorescent intensity (n = 3 technical replicates).

#### Characterization of PD1 expression levels CHO-K1 cells

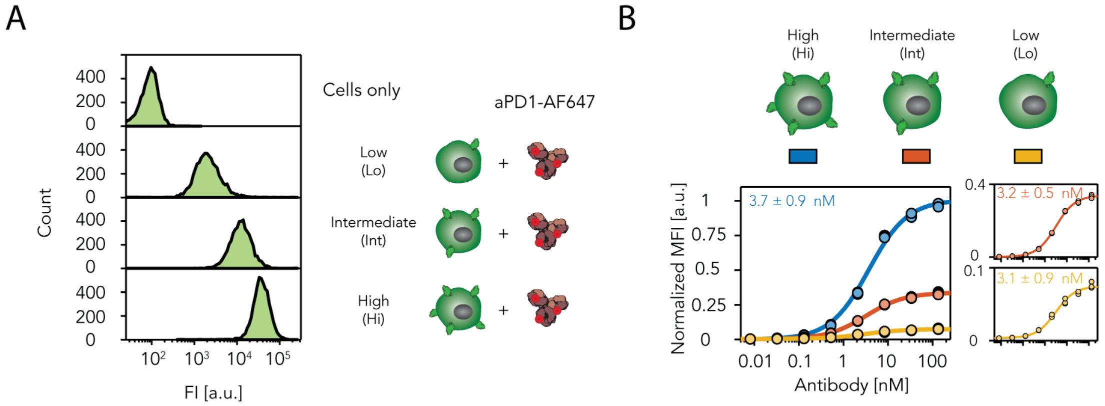

**Supplementary Figure 10 | Characterization of CHO-K1 cells expressing different levels of PD1. (a)** Flow cytometric analysis of CHO-K1 cells expressing low, intermediate and high levels of PD1 incubated with 10 nM of AF647-aPD1. **(b)** Flow cytometric analysis of AF647-aPD1 titration to CHO-K1 cells expressing low, intermediate and high levels of PD1. The mean fluorescent intensities, corrected for AF67 labeling efficiency, were fitted to a non-cooperative Hill equation to extract the apparent dissociation constant. Fluorescent intensity levels decreased as a function of expression levels while the $K_{d,app}$ was similar for all expression levels. Individual data points represent the normalized mean fluorescent intensity of 2000 gated single-cell events (n = 3 technical replicates).

##### DNA labeling of Cetuximab and Trastuzumab

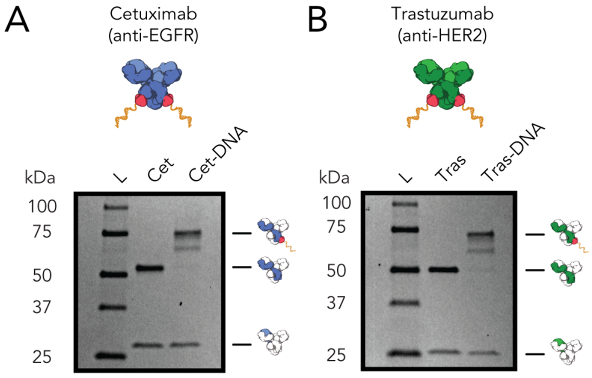

**Supplementary Figure 11 | SDS-PAGE analysis of an antibody protein G-DNA conjugation for Cetuximab and Trastuzumab.** Image of a reduced SDS-PAGE gel (4-20% polyacrylamide, stained for proteins with Coomassie Brilliant Blue) of the conjugation process for protein G-DNA to **(a)** Cetuximab and **(b)** Trastuzumab. Protein G-DNA (10 molar equivalents) conjugation to the Fc region of the antibody for 1h at 4 °C using 365 nm illumination results in a gel shift of the band corresponding to the heavy chain of the antibody. Since pG-DNA conjugates were not purified before coupling, a second band just below the final product is apparent which represents a heavy chain coupled to pG without a DNA handle. Gel densitometry as shown in **Supplementary Figure 2** was used to determine the concentration of each DNA conjugated antibody. Label: L, reference protein ladder.

##### Optimization of DNA nanostructure determinants for receptor binding efficiency

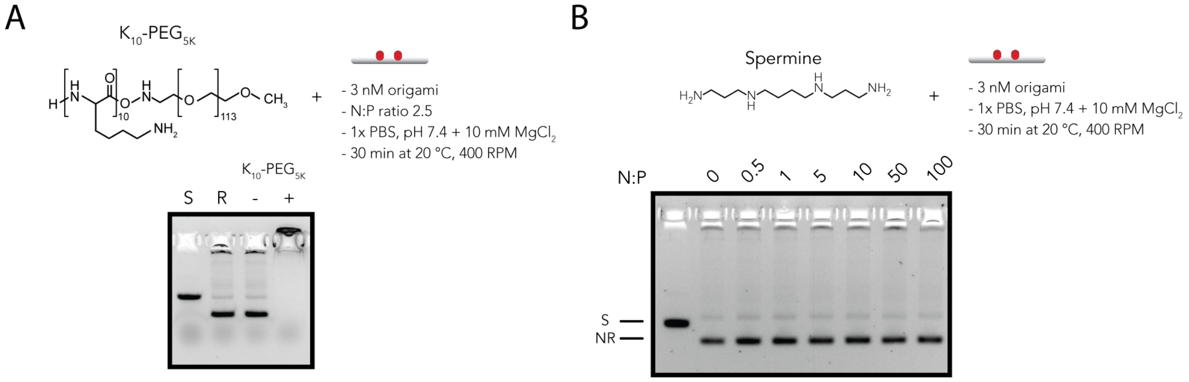

**Supplementary Figure 12 | Coating of DNA nanorods with different polyamines. (a)** DNA nanorods were incubated for 30 min with a **(a)** polyethylene glycol-oligolysine co-polymer ($K_{10}$-$PEG_{5K}$) using a N:P ratio (nitrogen in amines to phosphorus in DNA) of 2.5 or **(b)** spermine using different N:P ratio's. Agarose gel electrophoresis (1.5% agarose, stained for DNA with SYBR safe) was used to confirm coating and monitor formation of aggregation. For $K_{10}$-$PEG_{5K}$ it is expected that the coated nanostructure remains in the well due to the size of the PEG polymers.[4] Labels: S, scaffold; R, reference nanorod; NR, DNA nanorod.

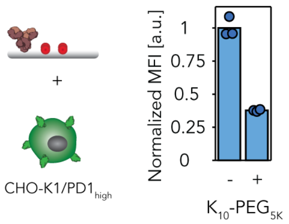

**Supplementary Figure 13 | Flow cytometric analysis of $K_{10}$-$PEG_{5K}$ coated aPD1-functionalized DNA nanorods binding to CHO-$K1_{PD1\text{-}high}$ cells.** Flow cytometric analysis of CHO-$K1_{PD1\text{-}high}$ cells that were incubated for 1h with 20 nM uncoated aPD1-nanorods or 20 nM aPD1-nanorods coated with $K_{10}$-$PEG_{5K}$ using a N:P ratio of 2.5. Individual data points represent the normalized mean fluorescent intensity of 2000 gated single-cell events (n = 3 technical replicates).

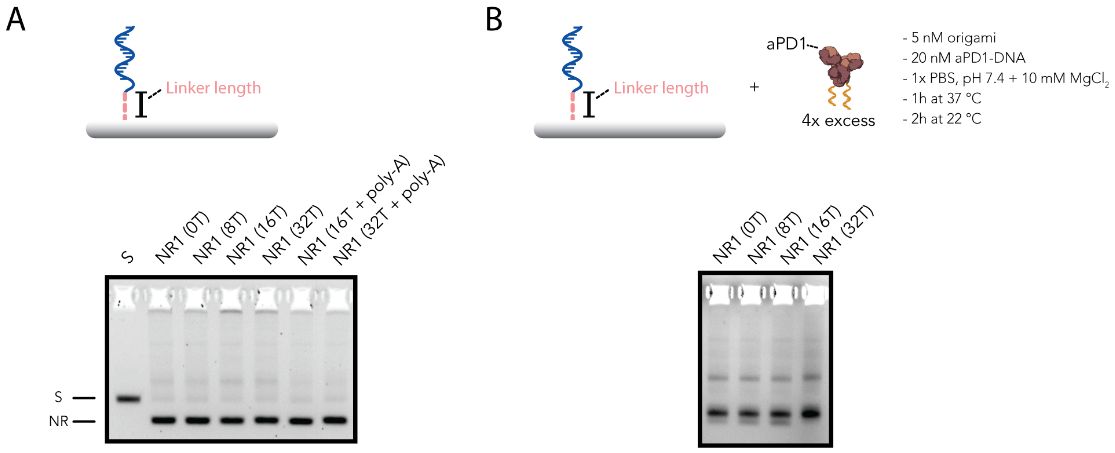

**Supplementary Figure 14 | Structural integrity of DNA nanorods self-assembled with different staple linker lengths. (a)** Image of an agarose gel (1.5% agarose, stained for DNA with SYBR safe) of DNA nanorods self-assembled with DNA handles that contained a 0, 8, 16 and 32-nucleotide (nt) single-stranded linker that separates the receptor from the NR surface. Additionally, DNA nanorods that contained a 16 and 32 nt linker were fortified with a complementary handle and loaded on the gel. **(b)** Image of an agarose gel of aPD1-DNA labeled DNA nanorods that contained a 0, 8, 16 and 32-nucleotide (nt) single-stranded linker. The similar incorporation efficiency of aPD1-DNA indicates that despite the difference in DNA handle length, the ssDNA handles were incorporated with a similar efficiency. The enhanced incorporation of aPD1-DNA conjugates on the nanorod functionalized with a 32-nt DNA handle is most likely a result of the increased flexibility of the DNA handle which facilitates incorporation of the antibody. Labels: S, scaffold; R, reference nanorod; NR, DNA nanorod.

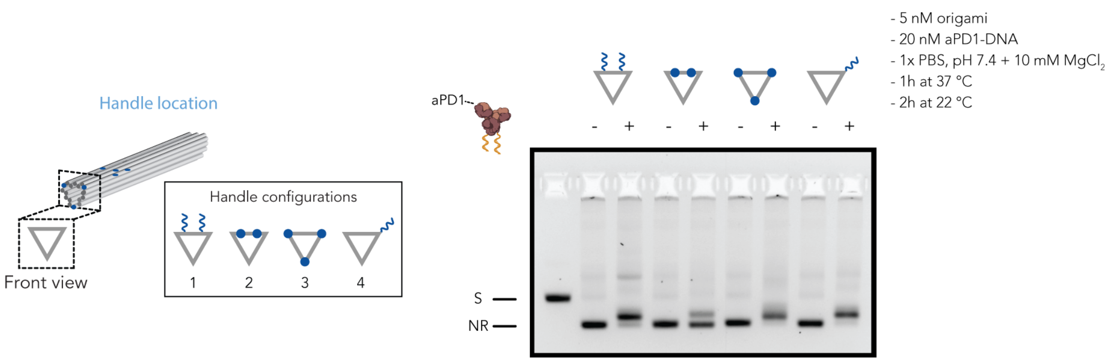

**Supplementary Figure 15 | Structural integrity of DNA nanorods self-assembled with DNA handles at defined locations.** Image of an agarose gel (1.5% agarose, stained for DNA with SYBR safe) of 4 configurations of DNA nanorods self-assembled with DNA handles at different locations. To assess the incorporation efficiency of DNA handles for each configuration, DNA nanorods were incubated with aPD1-DNA conjugates. Configuration 1, 3 and 4 showed similar incorporation efficiency of aPD1-DNA indicating similar availability of ssDNA handles. Configuration 2 displayed limited aPD1 incorporation, indicating that ssDNA handle incorporation was less efficient. Labels: S, scaffold; R; NR, DNA nanorod.

#### Design of multiple DNA nanostructures

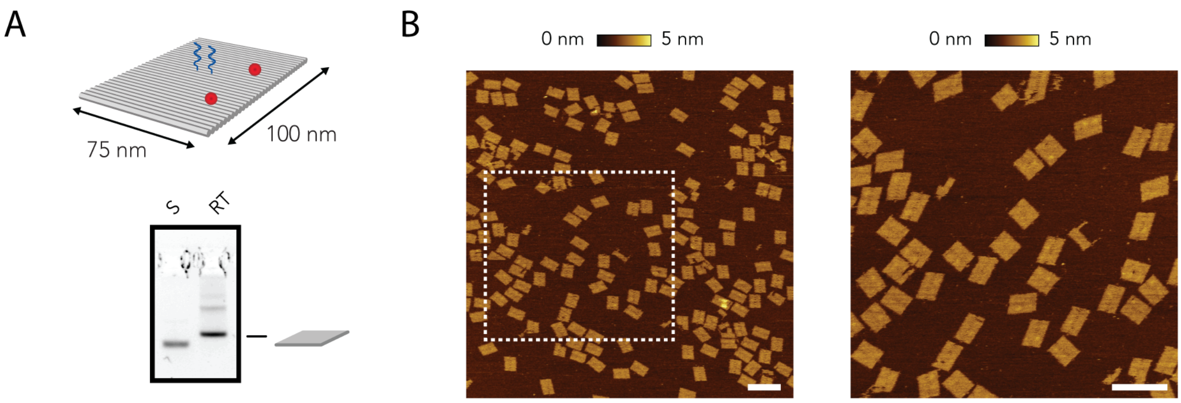

**Supplementary Figure 16 | Structural integrity and AFM analysis of DNA rectangle. (a)** Image of an agarose gel (1.5% agarose, stained for DNA with SYBR safe) of a self-assembled DNA origami rectangle (RT). **(b)** Topographic AFM image of the self-assembled DNA rectangle. Dashed rectangle indicates the region used for the detail image shown on the right. Scale bar, 200 nm.

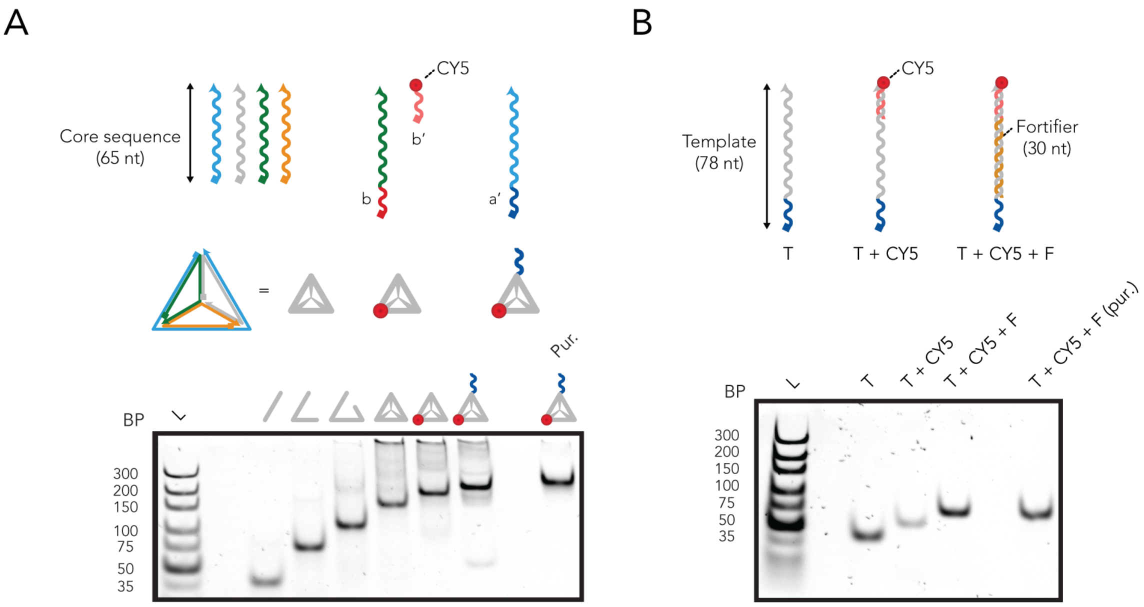

**Supplementary Figure 17 | Assembly of the DNA tetrahedron and the double stranded probe. (a)** Synthesis of the DNA tetrahedral nanostructure. The nanostructure is assembled from four single-stranded DNA strands. Two strands consist of a variable overhang (red and blue) and combinatorial core sequences (**Supplementary Table 6**). The complete stepwise formation and purification of the DNA tetrahedron is monitored with native polyacrylamide gel electrophoresis (PAGE) analysis (6%). **(b)** Image of a 6% native PAGE gel monitoring the self-assembly and purification of the double-stranded DNA probe (**Supplementary Table 5**). Labels: L, reference ladder; T, template; F, fortifier; Pur, purified.

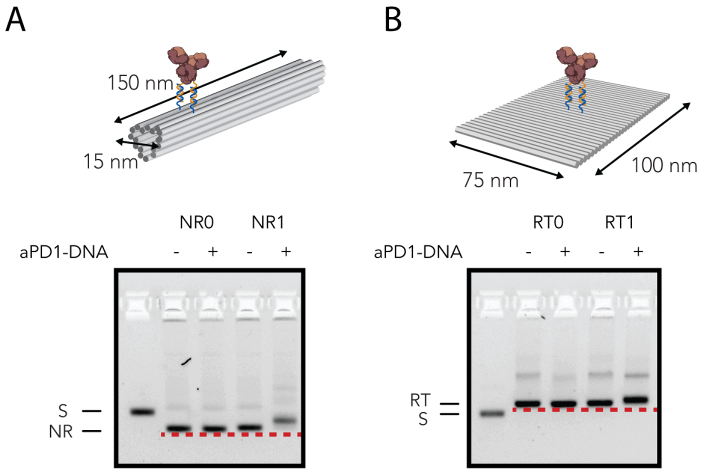

**Supplementary Figure 18 | Incorporation efficiency of aPD1-DNA conjugates on DNA nanorod and DNA rectangle.** Images of agarose gels (1.5% agarose, stained for DNA with SYBR safe) of **(a)** aPD1-functionalized nanorods or **(b)** aPD1-functionalized rectangle. No significant difference in incorporation efficiency of aPD1 is observed. Labels: S, scaffold; NR, DNA nanorod; RT, DNA rectangle.

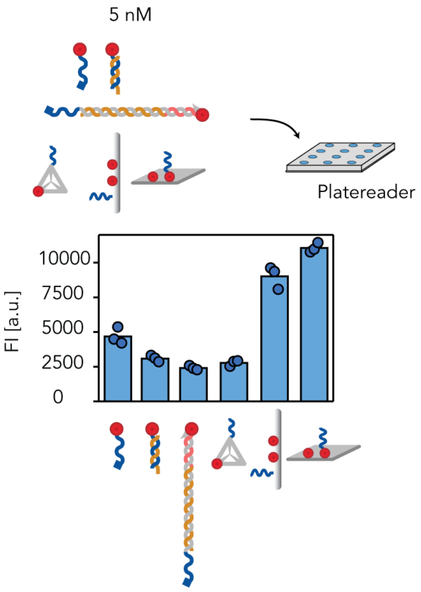

**Supplementary Figure 19 | Fluorescent intensity of constructed DNA nanostructures.** Fluorescent intensity measurements of 5 nM CY5-functionalized nanostructure. The fluorescent intensity of CY5-labeled imager strands (CY5-IM) was measured single-stranded as well as double-stranded to reflect binding of CY5-IM to the antibody-DNA conjugate. The fluorescent intensities of the double stranded CY5-labeled imager strand (IM), double-standed probe (dsP) and tetrahedron (Tet) are similar. In addition, CY5-functionalized DNA nanorods and rectangles also show comparable fluorescent intensities. The enhancement in fluorescent intensity when comparing single CY5 nanostructures to CY5-Rec and CY5-NR is ~3-fold and therefore not proportional. We rationalized that a correction factor would only have a minor impact on the absolute difference in fluorescent intensity between CY5-IM and CY5-NC observed for cellular binding assays (>10-fold) and therefore we directly displayed the mean fluorescent intensity measured by flow cytometry in **Figure 3** without a correction. Individual data points represent the fluorescent intensity (n = 3 technical replicates).

#### PD1/PD-L1 blocking assay characterization

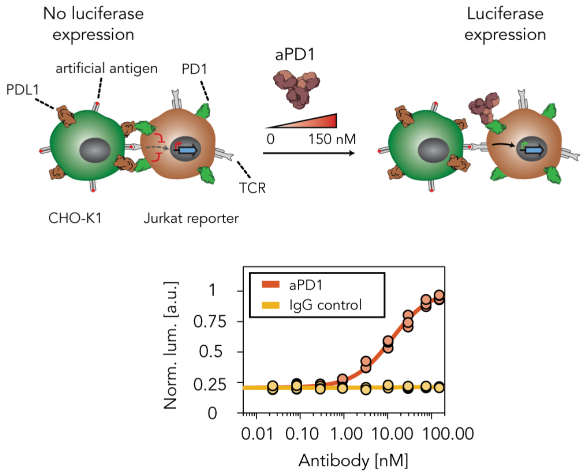

**Supplementary Figure 20 | Dose-dependent response of PD1 blocking.** Titration of aPD1 to a coculture of artificial antigen presenting (aAPC) CHO-K1$_{PD\text{-}L1/APC}$ cells that express the programmed death-ligand 1 (PD-L1) and Jurkat T$_{PD1/TCR}$ cells stably expressing PD1, T-cell receptors (TCRs) and nuclear factor of activated T cells (NFAT). Schematic of the cell assay to detect PD1 checkpoint inhibition using aPD1. Specifically, artificial antigen presenting (aAPC) CHO-K1 cells that express the programmed death-ligand 1 (PD-L1) were cocultured with a Jurkat T reporter cell line stably expressing PD1, T-cell receptors (TCRs) and nuclear factor of activated T cells (NFAT) induced luciferase. The luminescence was measured after 6h, normalized to the fitted maximum luminescence and fitted to a non-cooperative Hill equation. Individual data points represent the normalized luminescence of 3 technical replicates.

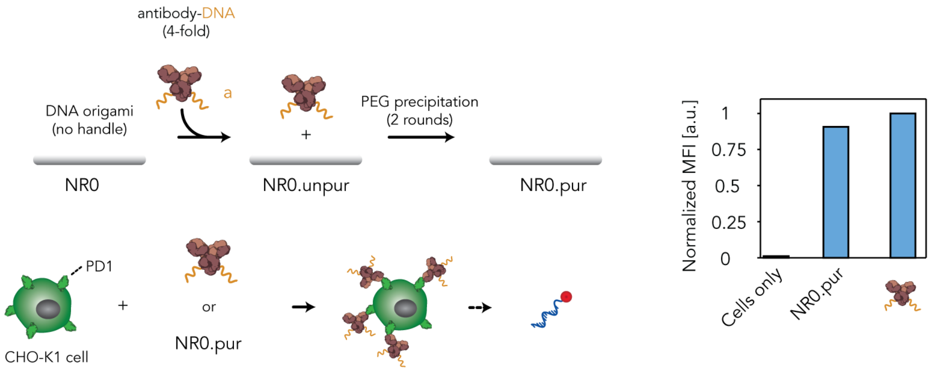

**Supplementary Figure 21 | Purification of aPD1-NR using PEG precipitation.** DNA nanorods self-assembled without ssDNA handles (NR0) were incubated with a 4-fold molar equivalent of aPD1-DNA for 1h at 37 °C and 2h at 22 °C (NR0.unpur). Subsequently, aPD1-DNA conjugates were removed using 2 rounds of PEG precipitation as described in the **Methods** section to obtain NR0.purifed which should be free of aPD1-DNA conjugates. CHO-K1$_{PD1\text{-}high}$ cells were incubated with 20 nM aPD1-DNA conjugates or 20 nM NR0.pur for 30 min at RT. After incubation, cells were spun down and reconstituted in 10 nM of CY5-labeled imager strands (CY5-IM) to fluorescently label aPD1-DNA conjugates for 30 min at RT. Flow cytometric analysis of the labeled cells is used to quantify aPD1 contamination. CHO-K1$_{PD1\text{-}high}$ cells incubated with NR0.pur display similar fluorescent intensity levels as cells incubated with aPD1-DNA conjugates, indicating that traces of free aPD1-DNA conjugates were still present in the sample after 2 rounds of PEG precipitation. Individual data points represent the normalized mean fluorescent intensity of 2000 gated single-cell events (n = 1).

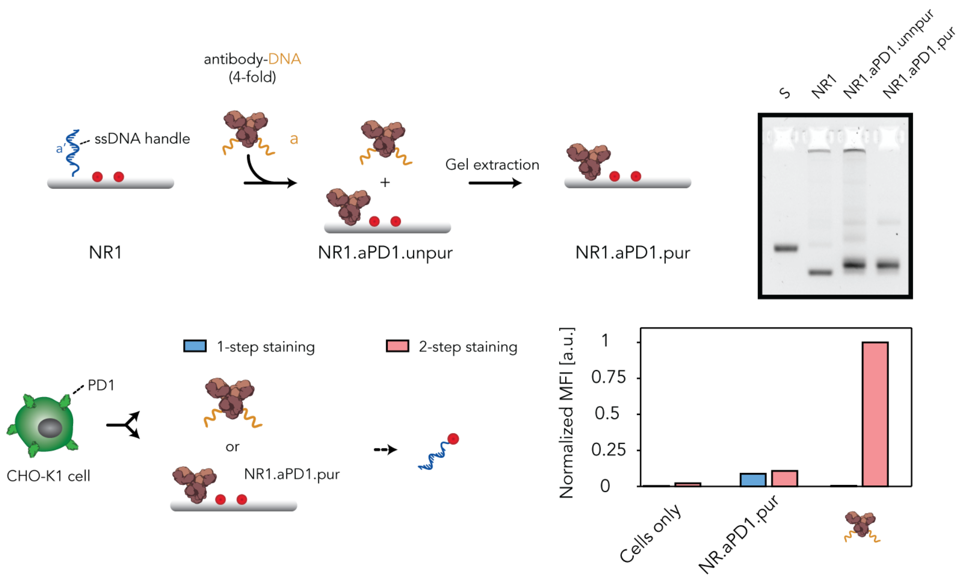

**Supplementary Figure 22 | Purification of aPD1-NR using agarose gel electrophoresis.** DNA nanorods self-assembled with ssDNA handles and two CY5 fluorophores (NR1) were incubated with a 4-fold molar equivalent of aPD1-DNA for 1h at 37 °C and 2h at 22 °C (NR0.unpur). Subsequently, aPD1-DNA conjugates were removed by loading the reaction mixture (NR1.aPD1.unpur) on an agarose gel (1.5%), running the gel for 90 min in an ice bath, and extracting the band corresponding to NR1.aPD1 to obtain NR1.aPD1.pur as described in the **Methods** section. The image of an agarose gel (1.5% agarose, stained for DNA with SYBR safe) confirms the structural integrity and aPD1-DNA incorporation after gel extraction. CHO-K1$_{PD1\text{-}high}$ cells were incubated with 20 nM aPD1-DNA conjugates or 20 nM NR1.aPD1.pur for 30 min at RT and fluorescent intensity levels were quantified using flow cytometry (blue bars). Subsequently, cells were spun down and reconstituted in 10 nM of CY5-labeled imager strands (CY5-IM) to fluorescently label aPD1-DNA conjugates for 30 min at RT (red bars). Flow cytometric analysis of the labeled cells is used to quantify aPD1 contamination. CHO-K1$_{PD1\text{-}high}$ cells incubated with NR1.aPD1.pur do not display an increased fluorescent intensity after incubating cells with CY5-IM, indicating the absence of aPD1-DNA conjugates and confirming appropriate purification. Individual data points represent the normalized mean fluorescent intensity of 2000 gated single-cell events (n = 1).

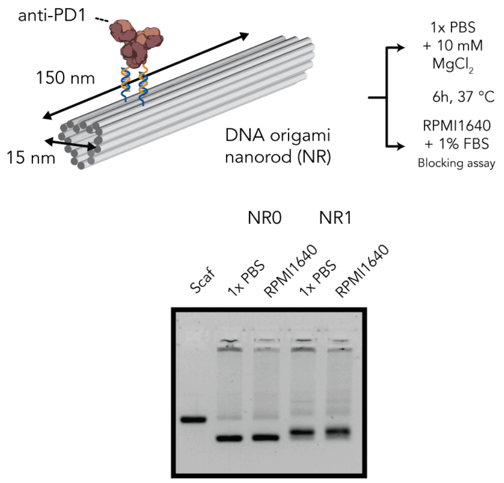

**Supplementary Figure 23 | Stability of aPD1-NR in culture medium.** Image of an agarose gel (1.5% agarose, stained for DNA with SYBR safe) to confirm the structural integrity of DNA nanorods after incubation for 6h in culture medium used for the PD1/PD-L1 blocking assay as shown in **Figure 5**. No significant degradation or loss of aPD1 is observed after 6h incubation in culture medium compared to 1x PBS and 10 mM $MgCl_2$.

#### Schematic of DNA origami nanostructures

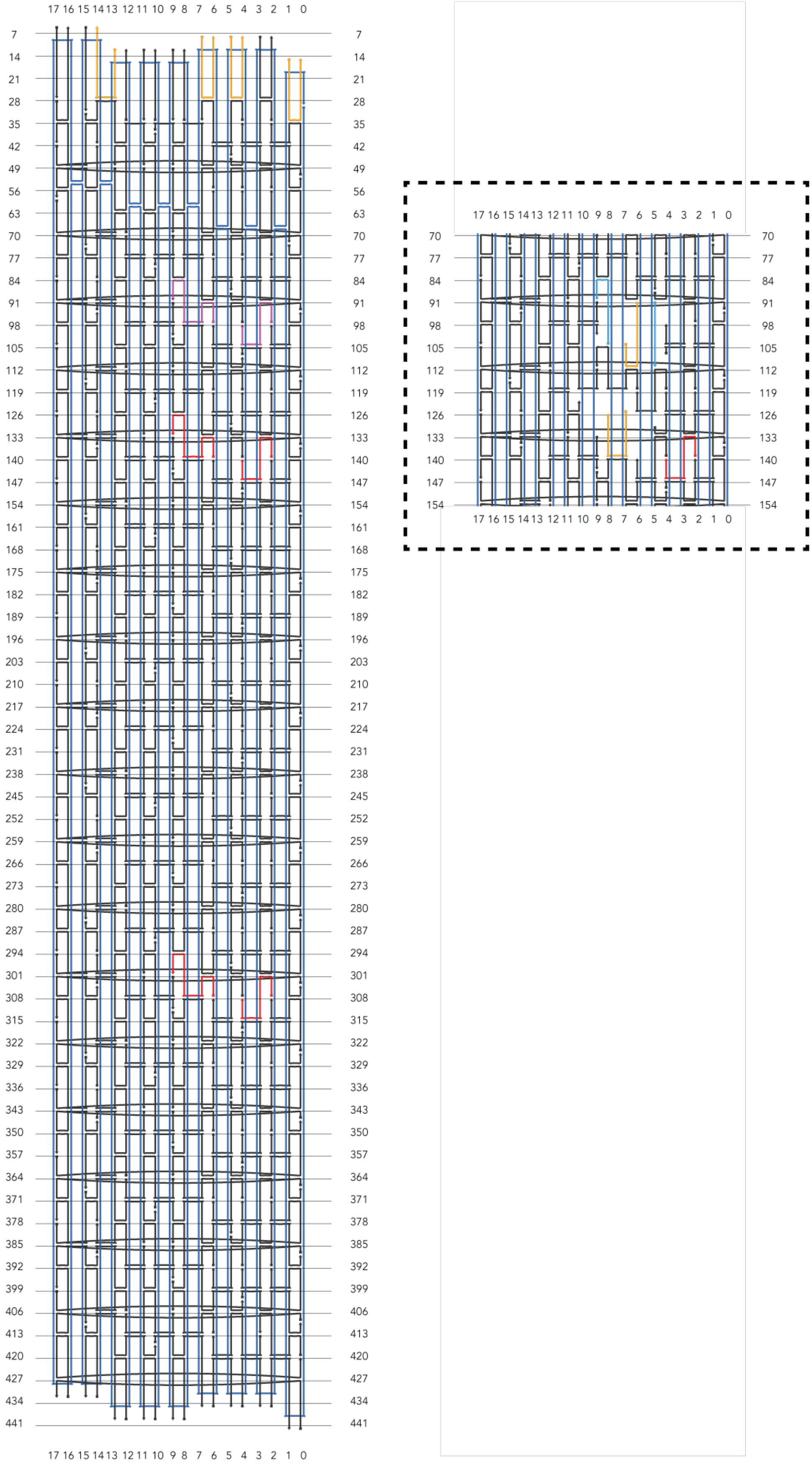

**Supplementary Figure 24 | Schematic overview of the DNA origami nanorod.** The scaffold strand is shown in light blue and unmodified core/edge staple strands in dark green (**Supplementary Table 8**). Staples that are commonly used for antibody incorporation are shown in purple (corresponding to staples a1 and a2, **Supplementary Table 3**). Staples that are used for incorporation of a fluorescent dye are highlighted in red (corresponding to staples f1-4, **Supplementary Table 3**). Staples indicated in yellow represent the staples used to design the different configurations as depicted in **Figure 3d** (corresponding to staples c5-8, **Supplementary Table 4**). Inset represents the schematic overview of the staples used to design configuration **4** of **Figure 3d** (corresponding to staples c3-4, **Supplementary Table 4**). The staples that are shown in cyan are additional core staples used to self-assemble configuration **4** of **Figure 3d** (corresponding to staples c1-2, **Supplementary Table 4**). Numbers on the top and bottom indicate helix number, while the numbers on the right and left indicate nucleotide positions.

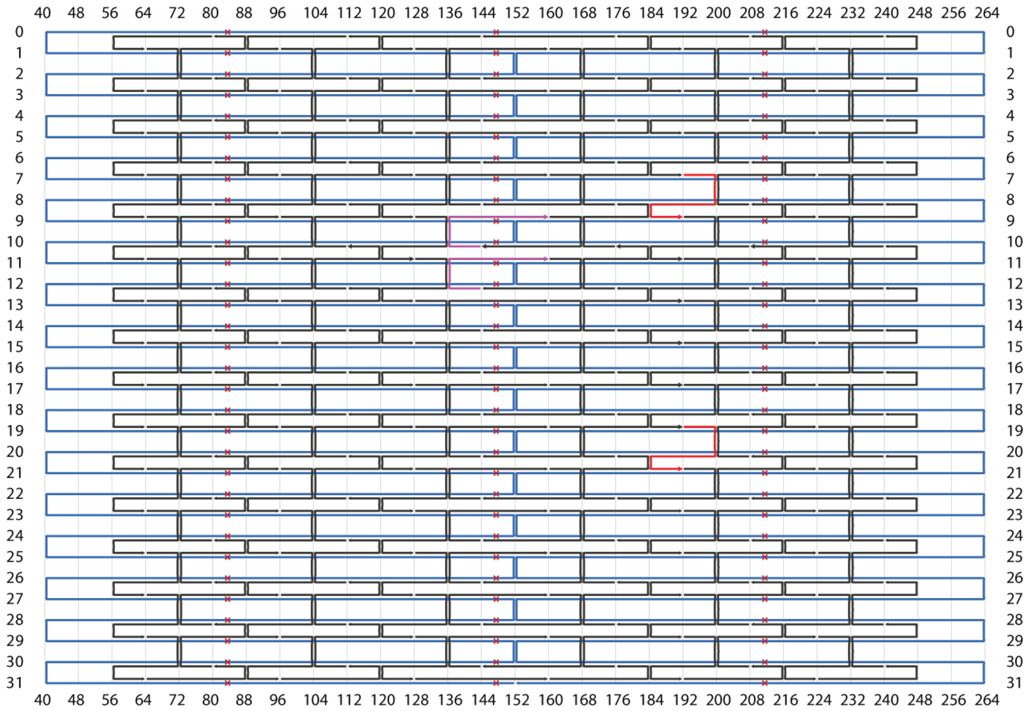

**Supplementary Figure 25 | Schematic overview of the DNA origami rectangle.** The scaffold strand is shown in light blue and unmodified core staple strands in dark green (**Supplementary Table 9**). Staples that are used for antibody incorporation are shown in purple (corresponding to staples r1-2, **Supplementary Table 7**). Staples that are used for incorporation of a fluorescent dye are highlighted in red (corresponding to staples r3-4, **Supplementary Table 7**). The red crosses indicate base-pair deletions to correct for global twist. Numbers on the top and bottom indicate nucleotide positions, while the numbers on the right and left indicate helix numbers.

### Supplementary tables

**Supplementary Table 1 | Anti-handle for protein G-DNA coupling and subsequent antibody labeling.** This oligo was used for NHS coupling to a maleimide and subsequent coupling to the N-terminal cysteine located in protein G. IDT modifications /3AmMO/ was used for amino modification at 3' end.

| ID | Sequence (5'to 3') |
|---|---|
| p1 | CCCTAGAGTGAGTCGTATGA/3AmMO/ |

**Supplementary Table 2 | Fluorescently labeled imagers**
To label DNA nanostructures and antibody-DNA conjugates, complementary imager (i) strands were used during the self-assembly of the DNA nanostructure (i1 & i2) or used in the cellular labeling process as a single stranded probe (i3).

| ID | Sequences (5'to 3') |
|---|---|
| i1 | **CAGTCAGTCAGTCAGTCAGT**TT/3AlexF647N/ |
| i2 | **CAGTCAGTCAGTCAGTCAGT**/3Cy5Sp/ |
| i3 | **TCATACGACTCACTCTAGGG**TT/3Cy5Sp/ |

**Supplementary Table 3 | Handle-extended staple strands antibody incorporation.**
To assemble DNA nanorods for anti-PD1 incorporation **(Figure 1** and **5)**, appropriate unmodified staples were replaced with handle-extended staple strands. Bold nucleotides refer to the extended handle sequence. Labels: a, antibody; f, fluorophore.

| ID | Unmodified staple ID | Used anti-handle | Sequences (5' to 3') |
|---|---|---|---|
| a1 | 186 | p1 | **TCATACGACTCACTCTAGGG**ATGCGCAAGAGTCTGGAGCAATAATGCC |
| a2 | 188 | p1 | **TCATACGACTCACTCTAGGG**AAAATCATGCTCCAAAGCGCGAAACAAACGCCACC |
| f1 | 190 | i1, i2 | **ACTGACTGACTGACTGACTG**GGAGAAGAAACTAGCATGTCAAATCACC |
| f2 | 192 | i1, i2 | **ACTGACTGACTGACTGACTG**AATGCTTCATAAGGAATACACTAAAACATTTCAGG |
| f3 | 204 | i1, i2 | **ACTGACTGACTGACTGACTG**TTCGCCAAATAATTCGCGTCTCTAAATC |
| f4 | 206 | i1 | **ACTGACTGACTGACTGACTG**GTAAGAGACGAGAATTTGCGGGATCGTCATTTTGC |

**Supplementary Table 4 | Handle-extended staple strands to tune design parameters.**

To assemble DNA nanorods with different handle extension lengths (e) or configurations (c) **(Figure 2)**, appropriate unmodified staples were replaced with handle-extended staple strands. Poly-adenine linker (l) strands l1 and l2 were used to produce double stranded staple extensions **Figure 2d**. Unless stated otherwise, all DNA nanorods where always assembled with handle-extended staple strand f2, f3 and i2. Underlined thymine nucleotides refer to the added spacer, whereas the bold nucleotides represent the extended handle sequence. To design configuration 2, 3 and 4 as shown in **Figure 2d** the following combinations of handle-extended strands were used: configuration **2**; c5 & c6, configuration **3**; c5, c7 & c8, configuration **4**; c1, c2, c3 and c4.

| ID | Unmodified staple ID | Used anti-handle | Sequences (5' to 3') |
|---|---|---|---|
| e1 | 186 | p1 | **TCATACGACTCACTCTAGGG**TTTTTTTTATGCGCAAGAGTCTGGAGCAA |
| e2 | 188 | p1 | **TCATACGACTCACTCTAGGG**TTTTTTTTAAAATCATGCTCCAAAGCGCGAAACAAA |
| e3 | 186 | p1 | **TCATACGACTCACTCTAGGG**TTTTTTTTTTTTTTTTATGCGCAAGAGTCTGGAGCAA |
| e4 | 188 | p1 | **TCATACGACTCACTCTAGGG**TTTTTTTTTTTTTTTTAAAATCATGCTCCAAAGCGCGAAACAAA |
| e5 | 186 | p1 | **TCATACGACTCACTCTAGGG**TTTTTTTTTTTTTTTTTTTTTTTTTTTTTTTTATGCGCAAGAGTCTGGAGCAA |
| e6 | 188 | p1 | **TCATACGACTCACTCTAGGG**TTTTTTTTTTTTTTTTTTTTTTTTTTTTTTTTAAAATCATGCTCCAAAGCGCGAAACAAA |
| l1 | - | e3, e4 | AAAAAAAAAAAAAAAA |
| l2 | - | e5, e6 | AAAAAAAAAAAAAAAAAAAAAAAAAAAAAAAA |
| c1 | 71 | - | TTATACCAAGCGCGAAACAAACGCCACC |
| c2 | 96 | - | AAAAAGATTAAGAGGAAGCCC |
| c3 | 187 | p1 | **TCATACGACTCACTCTAGGG**TTTTTTTTTTAGCCGGATGACCATAAATCAAAAATCA |
| c4* | 192 | p1 | **TCATACGACTCACTCTAGGG**TTTTTTTTCGGTCAATCATAAGGAATACACTAAAACA |
| c5 | 163 | p1 | **TCATACGACTCACTCTAGGG**TTTTTTTTCACTACGTGAACCATC |
| c6 | 165 | p1 | **TCATACGACTCACTCTAGGG**TTTTTTTTTGCGCCGCTACAGGGC |
| c7 | 166 | p1 | **TCATACGACTCACTCTAGGG**TTTTTTTTCAGGAACGGTACGCCA |
| c8 | 172 | p1 | **TCATACGACTCACTCTAGGG**TTTTTTTTTCACCAGTCACACTGAAAGCGTAAGA |

*) handle-extended staple c4 was used in combination with handle-extended staples f1 & f3 instead of f2 & f3.

**Supplementary Table 5 | Sequences of DNA strands used for the assembly of double stranded DNA probe (dsProbe).**

dsProbes (d) in **Figure 3** were assembled using 1 template strand that was 5' handle-extended for antibody-DNA binding and 3' extended for fluorophore incorporation. Furthermore, the template strand was fortified with spacer strand. Underlined thymine nucleotides refer to the added spacer, whereas the bold nucleotides represent the extended handle sequence.

| ID | Used anti-handle | Sequences (5' to 3') |
|---|---|---|
| d1 | p1 + i2 | **TCATACGACTCACTCTAGGG**TTTTTTTTAGTGAGTCCATGCTCAGGATTGCGAGTTAC**ACTGACTGACTGACTGACTG** |
| d2 | - | GTAACTCGCAATCCTGAGCATGGACTCACT |

**Supplementary Table 6 | Sequences of DNA strands used for the assembly of the tetrahedron.**

DNA tetrahedrons (t) in **Figure 3** were assembled using 4 core sequences that were handle-extended for antibody-DNA binding and fluorophore incorporation. In this table t5 is the handle-extended t1 and t6 represents handle-extended t3. Underlined thymine nucleotides refer to the added spacer, whereas the bold nucleotides represent the extended handle sequence.

| ID | Used anti-handle | Sequences (5' to 3') |
|---|---|---|
| t1 | - | TTCGAACATTCCTAAGTCTGAAATTTATCACCCGCCATAGTAGACGTATCACCAGGCAGTTGAGA |
| t2 | - | TTTTCAGACTTAGGAATGTTCGACATGCGAGGGTCCAATACCGACGATTACAGCTTGCTACACGA |
| t3 | - | TTCTACTATGGCGGGTGATAAAACGTGTAGCAAGCTGTAATCGACGGGAAGAGCATGCCCATCCA |
| t4 | - | TTCGGTATTGGACCCTCGCATGACTCAACTGCCTGGTGATACGAGGATGGGCATGCTCTTCCCGA |
| t5 | p1 | **TCATACGACTCACTCTAGGG**TTTTTTTTCGAACATTCCTAAGTCTGAAATTTATCACCCGCCATAGTAGACGTATCACCAGGCAGTTGAGA |
| t6 | i2 | **ACTGACTGACTGACTGACTG**TTCTACTATGGCGGGTGATAAAACGTGTAGCAAGCTGTAATCGACGGGAAGAGCATGCCCATCCA |

**Supplementary Table 7 | Handle-extended staple strands for the assembly of the DNA rectangle.**

To assemble DNA rectangles (r) for DNA-antibody binding in **Figure 3**, appropriate unmodified staples were replaced with handle-extended staple strands. Underlined thymine nucleotides refer to the added spacer, whereas the bold nucleotides represent the extended handle sequence.

| ID | Unmodified staple ID | Used anti-handle | Sequences (5' to 3') |
|---|---|---|---|
| r1 | 189 | p1 | **TCATACGACTCACTCTAGGG**TTTTTTTTAGCGTCCATAGTAAAATGTTTAGTTTATCCC |
| r2 | 190 | p1 | **TCATACGACTCACTCTAGGG**TTTTTTTTTCTTACCGGAAACAATGAAATAGCACTAACG |
| r3 | 191 | i2 | **ACTGACTGACTGACTGACTG**AGACAAAAGCAACATATAAAAGAAAAGTAAGC |
| r4 | 192 | i2 | **ACTGACTGACTGACTGACTG**AGACTACCCCCTTAGAATCCTTGAGATGAAAC |

**Supplementary Table 8 | Sequences of unmodified staple strands of the DNA nanorod.** Locations of the 5’ and 3’ end are indicated using the reference helix number used **in Supplementary Figure 24**, with the reference nucleotide position denoted in brackets.

| Staple ID | Location of 5' end | Location of 3' end | Sequence |
|---|---|---|---|
| 1 | 0[51] | 14[42] | AAGACACCGCCTAACTGGCGCGGTAAGCCAACAGAGAT |
| 2 | 0[93] | 14[94] | GGAGAAAAATAACAGTACTTGAAACAAG |
| 3 | 0[114] | 17[104] | GGTTGCTGAATGAATTACCTTTTTTAATGGACTAAAGC |
| 4 | 0[135] | 14[136] | CAACCCTCAAATTACATGTCAATAAGAA |
| 5 | 0[156] | 17[146] | TCGGTTGGCAAACATCAAGAAAACAAAATTATCAATAT |
| 6 | 0[177] | 14[178] | TGTAGGAATTGCAAAAGCTTTTTATAGA |
| 7 | 0[198] | 17[188] | TGCAAAATATTTATTCATTTCAATTACCTGAGAGGAAG |
| 8 | 0[219] | 14[220] | AGCACAACTAACAAAATAAATGCTCTGA |
| 9 | 0[240] | 17[230] | GCCTCAATAGGATTGCTTTGAATACCAAGTTATAGATT |
| 10 | 0[261] | 14[262] | AACGATTTAGGAAACAAAAACTTTAGCT |
| 11 | 0[282] | 17[272] | CTTACAAACAACAGTACCTTTTACATCGGGAAAGTATT |
| 12 | 0[303] | 14[304] | TTGATTAAATACGTCAGTAAATTTTAAA |
| 13 | 0[324] | 17[314] | CGGTTATTAATTGCGTAGATTTTCAGGTTTACCTTTGC |
| 14 | 0[345] | 14[346] | ATTGTAACATCGTAAAACGTTAAATTTC |
| 15 | 0[366] | 17[356] | GAAACAAAGAACCATATCAAAATTATTTGCATATCATT |
| 16 | 0[387] | 14[388] | CACGCGGAATATAATGGAGCCTGTGCCA |
| 17 | 0[408] | 17[398] | TCATGATTATTTGTTTGGATTATACTTCTGATATCATC |
| 18 | 0[419] | 3[412] | AGTGAGCCATACGAAACCGTGCATCTGCAATGGGA |
| 19 | 1[73] | 17[58] | TCAATTAAAGACGCTGAGTGTGAGTGAATAACCTTGCTTCTG |
| 20 | 2[34] | 4[35] | CCGATTTGAGAAAGGAAGGGAGCGCGTA |
| 21 | 2[55] | 2[56] | TCGGAACGAAAGGAGCGGGCGCTAGGAATGTAAAGCACTAAA |
| 22 | 2[76] | 4[77] | TTAGTGCAGGCTATCAGGTCATTTTTGA |
| 23 | 2[118] | 4[119] | CTTCTAAAATCGATGAACGGTCAACCGT |
| 24 | 2[160] | 4[161] | GGCCAGTGTACCCCGGTTGATGAGAAAG |
| 25 | 2[202] | 4[203] | GGTAACGTTGTATAAGCAAATATGCAAT |
| 26 | 2[244] | 4[245] | CCAGCTGTAAAATTCGCATTACAACGCA |
| 27 | 2[286] | 4[287] | CTGTTGGACCAATAGGAACGCCATTATG |
| 28 | 2[328] | 4[329] | CGGAAACCTGTAGCCAGCTTTATAAAGC |
| 29 | 2[370] | 4[371] | AAGATCGACCCGTCGGATTCTATAAATC |
| 30 | 3[413] | 6[413] | TAGGTCAAGGTGGCATCAATTCTGTTTAGCTATATACGAACT |
| 31 | 4[34] | 7[34] | ACCACCAGCGTACTATGGTTGAAACAGGAGGCCGAGAATCCT |
| 32 | 4[55] | 4[56] | GTAGCGGGAGCACGTATAACGTGCTTTCACGCGCTGGCAAGT |
| 33 | 4[76] | 6[77] | GAGATCTAGAAGCAAAGCGGAACCCTGA |
| 34 | 4[107] | 0[115] | AATACAAGAGGTGGTTGCCCGCTTCTAATCTATAG |
| 35 | 4[118] | 6[119] | TCTAGCTGAAAGACTTCAAATTTCAGAA |
| 36 | 4[149] | 0[157] | TCAATCATATGCCAAGCGATACCGACAGTGCGAAA |
| 37 | 4[160] | 6[161] | GCCGGAGAACCAGACCGGAAGGTCATAA |
| 38 | 4[191] | 0[199] | GTGAGGAAGACCAGGGTGTGGGCACGAATATGGTT |
| 39 | 4[202] | 6[203] | GCCTGAGTACCTTTAATTGCTAGTAAAA |
| 40 | 4[233] | 0[241] | TTTATTTTGTGCGAAAGACATAAATCATTTCCACC |
| 41 | 4[244] | 6[245] | AGGATAATGGCTTAGAGCTTAGCGAGAG |
| 42 | 4[275] | 0[283] | CTTTTTTTTAGAAGGGCATGAGTAAACAGGGTTTT |
| 43 | 4[286] | 6[287] | ACCCTGTATGTTTTAAATATGATCATAA |
| 44 | 4[317] | 0[325] | AAGGGCCTTCCAGGCAAATAAAGACGGAGGACGCG |
| 45 | 4[328] | 6[329] | CTCAGAGATTCCATATAACAGATAACGC |
| 46 | 4[359] | 0[367] | GCAAGTAACACACTCCATCATGGTCATAGCTTCGG |
| 47 | 4[370] | 6[371] | ATACAGGTTTAGTTTGACCATCAGTTGA |
| 48 | 4[401] | 0[409] | TAATGACCGTCAGTTTGACAATTCCACACAATAAC |
| 49 | 5[46] | 0[52] | GACTCACGCTAGAAAGCCCTAAAGTCAAGTTTTTTGGGGTCA |
| 50 | 5[88] | 0[94] | ATCGTAGCTATTGCCTGCGACTTAACAATGTCCCGCCAGTTT |
| 51 | 5[130] | 0[136] | CGTTGATATTAATCGTACCAGGGTCTCGCCCTGGAGTGAAAT |
| 52 | 5[172] | 0[178] | CTCAAAGGGTAATCAGACGTTGTACCATCTGTAAGCAAATCC |
| 53 | 5[214] | 0[220] | TTGATTTTAAATTTAAAAGGCGATTTGAATCGGCTGACTTGC |

| | | | |
|---|---|---|---|
| 54 | 5[256] | 0[262] | CTGTTTATTTAATTTTTTCTTCGCTCTGACCTCCTGGTGGGC |
| 55 | 5[298] | 0[304] | TAACCAAAAACATCAAATTCAGGCTACGTGGTGCTTGTCGTA |
| 56 | 5[340] | 0[346] | TTCTTAAGCACATCAACACCGCTTGTACCGAGCTCGAACTGC |
| 57 | 5[382] | 0[388] | TACACATCCACCGTGGGGTATCGGGTGTGAAATTGTTACGCT |
| 58 | 6[55] | 6[56] | TCAGAGCTAATCAGTGAGGGCCTGATAAAGTCCTCGTTAGAA |
| 59 | 6[76] | 9[69] | CTATTATATTGTGTGGAGATTTGTATCATAGTACC |
| 60 | 6[118] | 9[111] | AACGAGAAACGAGGTTGACCCCCAGCGACCTCAGA |
| 61 | 6[160] | 9[153] | ATATTCAAACTTTGCTAAAACGAAAGAGTAGGAAC |
| 62 | 6[202] | 9[195] | TGTTTAGATAGGCTTTAAACGGGTAAAAAAACTAC |
| 63 | 6[244] | 9[237] | GCTTTTGCGGATATGAGGCTTTGAGGACTAGCGTA |
| 64 | 6[286] | 9[279] | CCCTCGTCAGTGAAGCAGCGAAAGACAGTAAATGA |
| 65 | 6[328] | 9[321] | CAAAAGGAGTAAATTGCAGGGAGTTAAACAGTTTC |
| 66 | 6[370] | 9[363] | GATTTAGTGTGAATACCATCGCCCACGCTTGCGAA |
| 67 | 6[412] | 9[405] | AACGGAACAGTCAGTTAAACAGCTTGATAGGCTCC |
| 68 | 9[49] | 9[48] | GCCTGAGTAGAATTTCCCACCGAGTAAAAGAGTCTATCACTT |
| 69 | 9[70] | 12[70] | GCCACCCCACCGTACTCAGGATTAAAGGTGAATTAATTGACG |
| 70 | 9[91] | 11[90] | CTCAGAATATAGCCCGGAATACACCGAC |
| 71 | 9[102] | 4[108] | CACTTATACCTGTTACTTAGCCGGATGACCAGAAGCCCGATA |
| 72 | 9[112] | 12[112] | GCCACCACGTCGAGAGGGTTGAATTAGAGCCAGCACAAAAGG |
| 73 | 9[133] | 11[132] | GATAGCACAGTACCAGGCGGACAGTAGC |
| 74 | 9[144] | 4[150] | CAAGCAAAAGGAACCGAACTGACCTTGAATCCAAAGCGACAG |
| 75 | 9[154] | 12[154] | CCATGTATAGGATTAGCGGGGCAAGGCCGGAAACGCACAATC |
| 76 | 9[175] | 11[174] | TTCGTCAGAGACTCCTCAAGATGAAACC |
| 77 | 9[186] | 4[192] | TACTACGTAATACAGACCAGGCGCACTGGATTAGAGAGTAAT |
| 78 | 9[196] | 12[196] | AACGCCTACATGAAAGTATTACGTAATCAGTAGCGATAAAAG |
| 79 | 9[217] | 11[216] | ACAGCCCTTCGGAACCTATTACAAGTTT |
| 80 | 9[228] | 4[234] | AGTTAAAGACATCTTGACAAGAACCAAAAGATTGCGGAAAAT |
| 81 | 9[238] | 12[238] | ACGATCTCAGTTAATGCCCCCACTGTAGCGCGTTTTGTTAGC |
| 82 | 9[259] | 11[258] | TTTCCAGGAGTAACAGTGCCCCATTTTC |
| 83 | 9[270] | 4[276] | TAGCATCGGACAAAGCTGCTCATTTTACCAGGCTCAACAATA |
| 84 | 9[280] | 12[280] | ATTTTCTTTTTAACGGGGTCATATTAGCGTTTGCCCCCAAAA |
| 85 | 9[301] | 11[300] | TAAACAACAGGAGTGTACTGGCATAATC |
| 86 | 9[312] | 4[318] | CAAGGCCGCTACACCAGAACGAGTAATTACGAAGTTTCCATA |
| 87 | 9[322] | 12[322] | AGCGGAGCATACATGGCTTTTCAGAGCCACCACCGGTTACCA |
| 88 | 9[343] | 11[342] | AACAACTATTTACCGTTCCAGCTCCCTC |
| 89 | 9[354] | 4[360] | GAAATAACCGTCAACTTTAATCATGAATACCGAGTAGACAAG |
| 90 | 9[364] | 12[364] | TAATAATAATGGAAAGCGCAGCAGAACCGCCACCCCCGAAGC |
| 91 | 9[385] | 11[384] | AATCTCCATAAATCCTCATTACACCACC |
| 92 | 9[396] | 4[402] | AAAACCGATACTGGCTCATTATACCAACATTCAATAACCTAC |
| 93 | 9[406] | 12[406] | AAAAGGAGGCCTTGATATTCACAGAACCACCACCAATAATAA |
| 94 | 10[37] | 5[45] | GGTTAATAACGTCCATCGAGAAGTGTTTTTAGGGAGCTCTTT |
| 95 | 10[79] | 5[87] | TATTCAGAACGTACAACCGAAATCCGCGACCGGTCTTTTTGC |
| 96 | 10[121] | 5[129] | TGCCCCTCATCTCATCTCGCAGACGGTCAATTAAACAGATCG |
| 97 | 10[163] | 5[171] | GATCCGTAACCACCAACAAAGAGGACAGATGCGGAATCCAAA |
| 98 | 10[205] | 5[213] | GAAGTAGCATGTTTCCAGGCTGACCTTCATCGGGTAATCCTT |
| 99 | 10[247] | 5[255] | AAAAAAGTTTGGCTACATCATTACCCAAATCCAAAATAATTG |
| 100 | 10[289] | 5[297] | AAGGTATGGGACCCTCATAAGGCTTGCCCTGCAACACTCAAC |
| 101 | 10[331] | 5[339] | CGTTGAGAATTGAGGCTTGGGCTTGAGATGGCAGATACTTGA |
| 102 | 10[373] | 5[381] | CAGTTTTTCAGACAACATACCTTATGCGATTGATTCATTAGA |
| 103 | 11[49] | 11[48] | GAAATACCTACACAGGGAACTCAAACTATCGGCCTGCTCATG |
| 104 | 11[91] | 14[84] | TTGAGCCATTGAGGGAGGGAACCGGTATTTTTTAT |
| 105 | 11[133] | 14[126] | ACCATTAATGGTTTACCAGCGCTTGCGGGTATTAA |
| 106 | 11[175] | 14[168] | ATCGATACACGGAATAAGTTTTTTTGCACAATCAA |
| 107 | 11[217] | 14[210] | GCCTTTACATACATAAAGGTGTAACGAGAGAAAAA |
| 108 | 11[259] | 14[252] | GGTCATAAAGACTCCTTATTAAAACAGCGCAGAAC |
| 109 | 11[301] | 14[294] | AAAATCAAAACGCAATAATAATTTTGTTATTCTGT |
| 110 | 11[343] | 14[336] | AGAGCCGGTAAGCAGATAGCCAGAATAACCAGTAA |
| 111 | 11[385] | 14[378] | CTCAGAGGAAATAGCAATAGCAACTGAATTAACAA |
| 112 | 12[69] | 15[73] | GAAATTAAATCAGATCATTACCGCGCCCAAATCGTCGCTATTAATT |
| 113 | 12[111] | 15[115] | GCGACATGCGTTTTTCATCGAGAACAAGACATAGCGATAGCTTAGA |

| | | | |
|---|---|---|---|
| 114 | 12[153] | 15[157] | AATAGAATTAAATCTCCTTATCATTCCAGTGAATTTATCAAAATCA |
| 115 | 12[195] | 15[199] | AAACGCATATCCTGAATTTACGAGCATGACCTCCGGCTTAGGTTGG |
| 116 | 12[237] | 15[241] | AAACGTACTAATTTACAATAGATAAGTCGATGCAAATCCAATCGCA |
| 117 | 12[279] | 15[283] | GAACTGGCAATCCATAAACAACATGTTCTTCAAATATATTTTAGTT |
| 118 | 12[321] | 15[325] | GAAGGAAAAAATAGGTACCGACAAAAGGAATGGTTTGAAATACCGA |
| 119 | 12[363] | 15[367] | CCTTTTTAGCGCATTTAGGCAGAGGCATTAAGAATAAACACCGGAA |
| 120 | 12[405] | 15[409] | GAGCAAGTCAGAGGCTTAATTGAGAATCTTAGTATCATATGCGTTA |
| 121 | 13[49] | 13[48] | TTCTGTAGCAAGCATTTTTTGACGCTCAATCGTCTAGGGACA |
| 122 | 14[41] | 17[41] | AGAACCCTATTAGTCTTTAATATAGCCCTAAAACAGGCGGTC |
| 123 | 14[83] | 17[83] | TTTCATCTTTCCCTTAGAATCCATAAATCAATATAAGCCAGC |
| 124 | 14[93] | 9[101] | CCGTCTAAGAACGCGAGTCAACCGATTTGGGATATAAGCCGC |
| 125 | 14[125] | 17[125] | ACCAAGTAGACGCTGAGAAGATTAACAATTTCATTCCTCAAA |
| 126 | 14[135] | 9[143] | CGGGAGGTTTTGAAGCCAATTCATCCATTAGTTTTGCTAGCC |
| 127 | 14[167] | 17[167] | TAATCGGGTCTGAGAGACTACAAGATGATGAAACAAATCAAC |
| 128 | 14[177] | 9[185] | AACCCCAGCTACAATTTAAGACACGCAGCACAGAGGCTCCAG |
| 129 | 14[209] | 17[209] | TAATATCATATAACTATATGTCGCGCAGAGGCGAACTTTAGG |
| 130 | 14[219] | 9[227] | ACACGTCTTTCCAGAGCGAAAATAGCGTCAGTGCCTATTCAT |
| 131 | 14[251] | 17[251] | GCGCCTGCAAAGAACGCGAGATAACGGATTCGCCTATAATAC |
| 132 | 14[261] | 9[269] | AATCATATTATTTATCCCATGATTGCCCCCTGTGCCTTACGT |
| 133 | 14[293] | 17[293] | CCAGACGTTCATCTTCTGACCATGAATATACAGTAATTCGAC |
| 134 | 14[303] | 9[311] | GTATAACGTCAAAAATGACCGAGGCCGGAACGATGATACTTT |
| 135 | 14[335] | 17[335] | TAAGAGATGTGATAAATAAGGCAGAAATAAAGAAATTTTAAA |
| 136 | 14[345] | 9[353] | GAGCATAAAAACAGGGAAAGAAAACCACCCTTCTCTGAAAAG |
| 137 | 14[377] | 17[377] | CGCCAACTAATTACTAGAAAAAAGGGTTAGAACCTAACCACC |
| 138 | 14[387] | 9[395] | TATCACCCTGAACAAAGAAACAATCCGCCACCAAACAAAAAA |
| 139 | 15[32] | 10[38] | GGCTTCTGACCGACCAGTAATAAAGAAATGGGAAAAACTGCT |
| 140 | 15[74] | 10[80] | AATGTAGGAATATAGAAGGCTTATGGTAAATTCACCGTGGTG |
| 141 | 15[116] | 10[122] | TTAACCGCACAGCGAACCTCCCGACCAAAGAAAATCACTAAG |
| 142 | 15[158] | 10[164] | TAGCTGTCTTAAGATTAGTTGCTAATTTTGTTCACCAAGAAG |
| 143 | 15[200] | 10[206] | GTTCCATCCTAATCTTACCAACGCGCAACATACAGAATTTCT |
| 144 | 15[242] | 10[248] | AGATTTATCAGCCAGTTACAAAATCGCAGTATCATCGGGTAT |
| 145 | 15[284] | 10[290] | AATACGACAAAATAAGAAACGATTCGGAATAATCTTTTTAAT |
| 146 | 15[326] | 10[332] | CCGATATAAACAGCCTTTACAGAGGAACAAAGAACCGCTAAG |
| 147 | 15[368] | 10[374] | TCAATGTAATTAGACGGGAGAATTTATCTTATCAGAGCAAGC |
| 148 | 15[410] | 10[416] | TACAGTAGGGGTAATTGAGCGCTAAAGCCCAGAGCCGCGACG |
| 149 | 17[42] | 2[35] | AGTATTAGGCGAAAAACCGTCACCCAAAGGAGCCC |
| 150 | 17[59] | 1[72] | CAACAGTGCCAACGTGGACTCCAACGTCGAGGTGCCGAATTG |
| 151 | 17[84] | 2[77] | AGCAAATACAAGAGTCCACTACCTTATGAGTGTCC |
| 152 | 17[126] | 2[119] | TATCAAAAAGAATAGCCCGAGATTTACGGGATGTT |
| 153 | 17[168] | 2[161] | AGTTGAATTGATGGTGGTTCCGGCCCTGAAACGAC |
| 154 | 17[189] | 2[182] | GTTATCTCCCAGCAGGCGAAACTCGTCGTTTCCCA |
| 155 | 17[210] | 2[203] | AGCACTAAAGCGGTCCACGCTAGGGGCCTAAGTTG |
| 156 | 17[252] | 2[245] | ATTTGAGAGCTGATTGCCCTTTCCGAACTATTACG |
| 157 | 17[273] | 2[266] | AGACTTTTTCACCAGTGAGACTGGTGTAGATCGGT |
| 158 | 17[294] | 2[287] | AACTCGTGGCGCCAGGGTGGTCTTAAGCTGCGCAA |
| 159 | 17[315] | 2[308] | CCGAACGGGAGAGGCGGTTTGTACCTCGAGCGCCA |
| 160 | 17[336] | 2[329] | AGTTTGAAATGAATCGGCCAATCCCCGGCTGGTGC |
| 161 | 17[357] | 2[350] | TTGCGGAACCTGTCGTGCCAGTTCGTAAGCCAGCT |
| 162 | 17[378] | 2[371] | AGAAGGATGCCCGCTTTCCAGGTTTCCTCCTCAGG |
| 163 | 1[15] | 0[15] | AAAACACTACGTGAACCATCTATCAGGGCGATGGCCAAAA |
| 164 | 3[8] | 2[8] | AAAAAGCCGGCGAACGTGGCAGAGCTTGACGGGGAAAAAA |
| 165 | 5[8] | 4[8] | AAAATGCGCCGCTACAGGGCCACCCGCCGCGCTTAAAAAA |
| 166 | 7[8] | 6[8] | AAAACAGGAACGGTACGCCATTAAAGGGATTTTAGAAAAA |
| 167 | 8[34] | 8[12] | ACGCAAATTAACCGTTGTAAAAA |
| 168 | 9[12] | 9[34] | AAAAGCAATACTTCTTTGATTAG |
| 169 | 10[34] | 10[12] | AATATCCAGAACAATATTAAAAA |
| 170 | 11[12] | 11[34] | AAAACCGCCAGCCATTGCAACAG |
| 171 | 12[34] | 12[12] | ATTATTTACATTGGCAGATAAAA |
| 172 | 13[12] | 14[5] | AAAATCACCAGTCACACTGAAAGCGTAAGAATACGAAAA |
| 173 | 15[5] | 15[31] | AAAATGGCACAGACAATATTTTTGAAT |

| | | | |
|---|---|---|---|
| 174 | 17[5] | 17[27] | AAAAACCAGCAGAAGATAAAACA |
| 175 | 17[28] | 16[5] | GAGGTGATCGCCATTAAAAATACCGAACGAACCAAAA |
| 176 | 0[442] | 17[432] | AAAAAAGCCTGGGGTGTCAAAA |
| 177 | 2[435] | 1[442] | AAAATGGGCGCATCGTGCCGGAAGCATAAAGTGTAAAAA |
| 178 | 4[435] | 3[435] | AAAACGCGAGCTGAAACGTTGGTGTAGAAAAA |
| 179 | 6[435] | 5[435] | AAAATACGTTAATAAATTTCATTTGGGGAAAA |
| 180 | 8[439] | 9[439] | AAAAAGCTTGCTTTCGAGGTATTGTATCGGTTTATCAAAA |
| 181 | 10[415] | 7[435] | ATTGCCTTTAGAATTTCGACGTTGGGAAGAAAAATCAAAA |
| 182 | 10[440] | 11[440] | AAAAAGGTTGAGGCAGGTCACGCCAGCATTGACAGGAAAA |
| 183 | 12[439] | 13[439] | AAAACCACAAGAATTGAGTTATATCAGAGAGATAACAAAA |
| 184 | 14[432] | 15[432] | AAAAAAAGCCAACGCTCAACAAATTCTTACCAGTATAAAA |
| 185 | 16[432] | 0[420] | AAAAATCAATATAATCCTGACAGATGATGGCAATCCTAATG |
| 186 | 2[97] | 4[98] | ATGCGCAAGAGTCTGGAGCAATAATGCC |
| 187 | 4[97] | 6[98] | GGAGAGGAAAAAGATTAAGAGTAAATCA |
| 188 | 6[97] | 9[90] | AAAATCATGCTCCAAAGCGCGAAACAAACGCCACC |
| 189 | 17[105] | 2[98] | ATCACCTTGAGTGTTGTTCCAAAATAACTGAATTC |
| 190 | 2[139] | 4[140] | GGAGAAGAAACTAGCATGTCAAATCACC |
| 191 | 4[139] | 6[140] | ATCAATATTTAATTCGAGCTTCCCCTCA |
| 192 | 6[139] | 9[132] | AATGCTTCATAAGGAATACACTAAAACATTTCAGG |
| 193 | 17[147] | 2[140] | CTGGTCAGCAAAATCCCTTATACTCTATTTTCTCA |
| 194 | 2[181] | 4[182] | GTCACGAAAAGCCCCAAAAACTAGGTAA |
| 195 | 4[181] | 6[182] | AGATTCACAACAGGTCAGGATAGCGTCC |
| 196 | 6[181] | 9[174] | AATACTGAACGGTGTGCCACTACGAAGGACTGAGT |
| 197 | 2[223] | 4[224] | TGCTGCATTGTAAACGTTAATTAGAACC |
| 198 | 4[223] | 6[224] | CTCATATATAAGAGGTCATTTAGTTTTG |
| 199 | 6[223] | 9[216] | CCAGAGGAAGAGTATTTTTCATGAGGAATCCACAG |
| 200 | 17[231] | 2[224] | AGAGCCGTGGCCCTGAGAGAGGCATTTCGGGGATG |
| 201 | 2[265] | 4[266] | GCGGGCCGTTAAATCAGCTCATTGCGGG |
| 202 | 4[265] | 6[266] | AGAAGCCAATATAATGCTGTAACGACGA |
| 203 | 6[265] | 9[258] | TAAAAACAACGTAAACGAGGGTAGCAACTGTCGTC |
| 204 | 2[307] | 4[308] | TTCGCCAAATAATTCGCGTCTCTAAATC |
| 205 | 4[307] | 6[308] | GGTTGTAAGTACGGTGTCTGGAGGCATA |
| 206 | 6[307] | 9[300] | GTAAGAGACGAGAATTTGCGGGATCGTCATTTTGC |
| 207 | 2[349] | 4[350] | TTCCGGCATTAAATGTGAGCGAAGAATT |
| 208 | 4[349] | 6[350] | AGCAAAACCAATTCTGCGAACACATTCA |
| 209 | 6[349] | 9[342] | ACTAATGTTTAATTATATATTCGGTCGCAGAAAGG |
| 210 | 2[391] | 4[392] | GACGACAAACAAACGGCGGATTAGTAGT |
| 211 | 4[391] | 6[392] | AGCATTAATTTCGCAAATGGTATTACAG |
| 212 | 6[391] | 9[384] | GTAGAAATTAAGAAGTTGCGCCGACAATCGTTGAA |
| 213 | 17[399] | 2[392] | ATATTCCCATTAATTGCGTTGTCCGCTCAGGGGAC |

**Supplementary table 9 | Sequences of unmodified staple strands of the DNA rectangle.** Locations of the 5’ and 3’ end are indicated using the reference helix number used in **Supplementary Figure 25**, with the reference nucleotide position denoted in brackets.

| Staple ID | Location of 5'end | Location of 3'end | Sequence |
|---|---|---|---|
| 1 | 0[79] | 1[63] | ACTGAGTTTCGTCACCAGTACAAATCATAGTT |
| 2 | 0[111] | 1[95] | GCAAGCCCAATAGGAACCCATGTACGTCTTTC |
| 3 | 0[143] | 1[127] | CCTCAGAGCCACCACCCTCATTTTGTATGGGA |
| 4 | 0[175] | 0[144] | CCCTCAGAACCGCCACCCTCAGAACCGCCAC |
| 5 | 0[207] | 1[191] | TATCACCGTACTCAGGAGGTTTAGATTATTCT |
| 6 | 0[239] | 1[223] | GGGTTGATATAAGTATAGCCCGGATGAGACTC |
| 7 | 1[64] | 3[63] | AGCGTAACAAAAGGCTCCAAAAGGTTCGAGGT |
| 8 | 1[96] | 3[95] | CAGACGTTAATAATTTTTTCACGTCGATAGTT |
| 9 | 1[128] | 3[127] | TTTTGCTAAATAGAAAGGAACAACGCCCACGC |
| 10 | 1[160] | 2[144] | TGCCCCCTAACAGTGCCCGTATAATTTCAGC |
| 11 | 1[192] | 3[191] | GAAACATGTAATAAGTTTTAACGGAGGTTGAG |
| 12 | 1[224] | 3[223] | CTCAAGAGCATGGCTTTTGATGATTATTCACA |
| 13 | 2[79] | 0[80] | CTCCAAAAGATCTAAAGTTTTGTCCGTAAC |
| 14 | 2[111] | 0[112] | TTGCGAATAGTAAATGAATTTTCTCAGGGATA |
| 15 | 2[143] | 1[159] | GGAGTGAGAACAACTTTCAACAGACAGTTAA |
| 16 | 2[175] | 0[176] | CCTTGAGTGCCTATTTCGGAACCTTACCGCCA |
| 17 | 2[207] | 0[208] | TGTACTGGAAAGTATTAAGAGGCATAGGTG |
| 18 | 2[239] | 0[240] | GCGTCATAAAGGATTAGGATTAGCCGTCGAGA |
| 19 | 3[64] | 5[63] | GAATTTCTCAACGGCTACAGAGGCTTCCATTA |
| 20 | 3[96] | 5[95] | GCGCCGACGCAGCGAAAGACAGCACTACGAAG |
| 21 | 3[128] | 5[127] | ATAACCGATAAAGGCCGCTTTTGCAAAAGAAT |
| 22 | 3[160] | 4[144] | AGAGCCGCAGAGCCGCCACCAGAAAGGCTTG |
| 23 | 3[192] | 5[191] | GCAGGTCATCAGAACCGCCACCCTTTTGCCTT |
| 24 | 3[224] | 5[223] | AACAAATAACCGGAACCGCCTCCCTCATCGGC |
| 25 | 4[79] | 2[80] | GAGGGTAGTAAACAGCTTGATACTGAAAAT |
| 26 | 4[111] | 2[112] | CACCCTCAAATGACAACAACCATCTAAAGGAA |
| 27 | 4[143] | 3[159] | CAGGGAGTTATATTCGGTCGCTGCCACCACC |
| 28 | 4[175] | 2[176] | CCACCCTCCGCCAGCATTGACAGGGGTCAGTG |
| 29 | 4[207] | 2[208] | CGCCACCCGACGATTGGCCTTGAACAGGAG |
| 30 | 4[239] | 2[240] | GAGCCACCAATCCTCATTAAAGCCTCCAGTAA |
| 31 | 5[64] | 7[63] | AACGGGTACGACCTGCTCCATGTTATAAGGGA |
| 32 | 5[96] | 7[95] | GCACCAACATCATCGCCTGATAAAAGGACAGA |
| 33 | 5[128] | 7[127] | ACACTAAACAAGCGCGAAACAAAGTAGGCTGG |
| 34 | 5[160] | 6[144] | GTAATCAGAATGAAACCATCGATACCCCAGC |
| 35 | 5[192] | 7[191] | TAGCGTCACCATTACCATTAGCAAAGCGCCAA |
| 36 | 5[224] | 7[223] | ATTTTCGGGGAATTAGAGCCAGCAGATTGAGG |
| 37 | 6[79] | 4[80] | GAAATCCGAAATACGTAATGCCATCGGAAC |
| 38 | 6[111] | 4[112] | AGATTTGTCTAAAACGAAAGAGGCGGGATCGT |
| 39 | 6[143] | 5[159] | GATTATACACACTCATCTTTGACGCAGCACC |
| 40 | 6[175] | 4[176] | ACGTCACCTAGCGACAGAATCAAGCAGAGCCA |
| 41 | 6[207] | 4[208] | CAGTAGCAGACTGTAGCGCGTTTTCAGAGC |
| 42 | 6[239] | 4[240] | GCCATTTGTCATAGCCCCCTTATTCGGAACCA |
| 43 | 7[64] | 9[63] | ACCGAACTAAACACCAGAACGAGTCTTTAATC |
| 44 | 7[96] | 9[95] | TGAACGGTTCATTCAGTGAATAAGTTAAGAAC |
| 45 | 7[128] | 9[127] | CTGACCTTATTCATTACCCAAATCTTGGGAAG |
| 46 | 7[160] | 8[144] | AATAGAAAGAATAAGTTTATTTTGTGACAAG |
| 47 | 7[224] | 9[223] | GAGGGAAGAGCAAACGTAGAAAATAAGGAAAC |
| 48 | 8[79] | 6[80] | CTGACGAGGACCAACTTTGAAAGTTGTGTC |
| 49 | 8[111] | 6[112] | AAAGCTGCGTACAGACCAGGCGCATACAACGG |
| 50 | 8[143] | 7[159] | AACCGGATCATCAAGAGTAATCTTCACAATC |
| 51 | 8[175] | 6[176] | ACACCACGATTCATATGGTTTACCGGCCGGAA |
| 52 | 8[207] | 6[208] | TAAAGGTGGGGCGACATTCAACCAAATCAC |
| 53 | 8[239] | 6[240] | AGTATGTTGTAAATATTGACGGAACGACTTGA |

| | | | |
|---|---|---|---|
| 54 | 9[64] | 11[63] | ATTGTGAAACATAACGCCAAAAGGTAACCCTC |
| 55 | 9[96] | 11[95] | TGGCTCATTTTAGGAATACCACATCAAAATAG |
| 56 | 9[128] | 11[127] | AAAAATCTTTATTACAGGTAGAAATTGCCAGA |
| 57 | 9[192] | 11[191] | AGATAGCCAAGAATTGAGTTAAGCGTTTAACG |
| 58 | 9[224] | 11[223] | CGAGGAAAATTGAGCGCTAATATCTACAGAGA |
| 59 | 10[79] | 8[80] | ATGCAGATTTACCTTATGCGATTGCTTGCC |
| 60 | 10[111] | 8[112] | AGTTGAGATATACCAGTCAGGACGAACGTAAC |
| 61 | 10[143] | 9[159] | GAACAACAACGTTAATAAAACGAAATAGCTA |
| 62 | 10[175] | 8[176] | AAGAGCAAAAGCCCTTTTTAAGAAACGCAAAG |
| 63 | 10[207] | 8[208] | TAACCCACGAACAAAGTTACCAGACATACA |
| 64 | 10[239] | 8[240] | AGAGGGTACGCAATAATAACGGAATATTACGC |
| 65 | 11[64] | 13[63] | GTTTACCAACGAGAATGACCATAAAGTCAGAA |
| 66 | 11[96] | 13[95] | CGAGAGGCTCCCCCTCAAATGCTTATTAAGAG |
| 67 | 11[128] | 13[127] | GGGGGTAAATACTGCGGAATCGTCGCGTTTTA |
| 68 | 11[160] | 12[144] | AATCCAAAATAAACAGCCATATTAACTGGAT |
| 69 | 11[192] | 13[191] | TCAAAAATGTCTTTCCAGAGCCTAAGGCTTAT |
| 70 | 11[224] | 13[223] | GAATAACATTTATCCTGAATCTTAGTTTTAGC |
| 71 | 12[79] | 10[80] | TTCAGAAAGACGACGATAAAAACTCAACTA |
| 72 | 12[111] | 10[112] | TCATTGAATTTTGCAAAAGAAGTTGATTCATC |
| 73 | 12[175] | 10[176] | GTTACAAATAAGAAACGATTTTTTCCAATAAT |
| 74 | 12[207] | 10[208] | TAACGAGCGAAAATAGCAGCCTTAGAGAGA |
| 75 | 12[239] | 10[240] | GCTACAATTAAAAACAGGGAAGCGACAAAGTC |
| 76 | 13[64] | 15[63] | GCAAAGCGTGGCTTAGAGCTTAATTAAATATG |
| 77 | 13[96] | 15[95] | GAAGCCCGGCTCCTTTTGATAAGAGTTTCATT |
| 78 | 13[128] | 15[127] | ATTCGAGCCAACAGGTCAGGATTACTGCGAAC |
| 79 | 13[160] | 14[144] | AATAGCAAATCGTAGGAATCATTAACCGGAA |
| 80 | 13[192] | 15[191] | CCGGTATTTCATCGAGAACAAGCAACGCGCCT |
| 81 | 13[224] | 15[223] | GAACCTCCCCAAGAACGGGTATTATGAACAAG |
| 82 | 14[79] | 12[80] | TTTGCGGAGATTGCATCAAAAAGTAAACAG |
| 83 | 14[111] | 12[112] | CTTTAATTAAAGACTTCAAATATCATAAATAT |
| 84 | 14[143] | 13[159] | GCAAACTCTTCAAAGCGAACCAGCCGCGCCC |
| 85 | 14[175] | 12[176] | TTATTTTCGCAAATCAGATATAGAATTTGCCA |
| 86 | 14[207] | 12[208] | TACCGCACCTAAGAACGCGAGGCCCAACGC |
| 87 | 14[239] | 12[240] | TTATCATTCGACTTGCGGGAGGTTTGCACCCA |
| 88 | 15[64] | 17[63] | CAACTAAACTACTAATAGTAGTAGAAAGAATT |
| 89 | 15[96] | 17[95] | CCATATAATGGGGCGCGAGCTGAACAGAGCAT |
| 90 | 15[128] | 17[127] | GAGTAGATTGGTCAATAACCTGTTACATTATG |
| 91 | 15[160] | 16[144] | AACAACATAATTCTGTCCAGACGAGATACAT |
| 92 | 15[192] | 17[191] | GTTTATCAAGAGAATATAAAGTACAAAAGCCT |
| 93 | 15[224] | 17[223] | AAAAATAATTTAGGCAGAGGCATTAAATTCTT |
| 94 | 16[79] | 14[80] | CATCAATTGTACGGTGTCTGGAAGGTCATT |
| 95 | 16[111] | 14[112] | TTTTCATTCAGTTGATTCCCAATTGAGAGTAC |
| 96 | 16[143] | 15[159] | TTCGCAAATTAGTTTGACCATTACGACAATA |
| 97 | 16[175] | 14[176] | GGTAAAGTGTTCAGCTAATGCAGAAGCCGTTT |
| 98 | 16[207] | 14[208] | CAGTAATAACAATAGATAAGTCCAACCAAG |
| 99 | 16[239] | 14[240] | ACATGTAATATCCCATCCTAATTTGTCTTTCC |
| 100 | 17[64] | 19[63] | AGCAAAATGTAAAGATTCAAAAGGCAATATGA |
| 101 | 17[96] | 19[95] | AAAGCTAAATTTTAAATGCAATGCATTAATGC |
| 102 | 17[128] | 19[127] | ACCCTGTACGCAAGGATAAAAATTGATCTACA |
| 103 | 17[160] | 18[144] | AACACCGGTAAATAAGGCGTTAAAAAGCCTT |
| 104 | 17[192] | 19[191] | GTTTAGTAAAATTTAATGGTTTGAGGTCTGAG |
| 105 | 17[224] | 19[223] | ACCAGTATATATATTTTAGTTAATTTAGGTTG |
| 106 | 18[79] | 16[80] | ATGTGTAGTAAGCAATAAAGCCTAAGGTGG |
| 107 | 18[111] | 16[112] | CCTCATATATCGGTTGTACCAAAATAGCTATA |
| 108 | 18[143] | 17[159] | TATTTCAAATACTTTTGCGGGAGTAAGAATA |
| 109 | 18[175] | 16[176] | CCGTGTGAAATCATAATTACTAGACGACAAAA |
| 110 | 18[207] | 16[208] | TCTGACCTTCATATGCGTTATACTTCGAGC |
| 111 | 18[239] | 16[240] | TTTTTCAAAAAGCCAACGCTCAACCAACGCCA |
| 112 | 19[64] | 21[63] | TATTCAACTCAGAAAAGCCCCAAATGTAAACG |
| 113 | 19[96] | 21[95] | CGGAGAGGGCATGTCAATCATATGTTAAATTT |

| | | | |
|---|---|---|---|
| 114 | 19[128] | 21[127] | AAGGCTATAAGAGAATCGATGAACCCAATAGG |
| 115 | 19[160] | 20[144] | AATAGTGATAGATTAAGACGCTGAGAGTCTG |
| 116 | 19[224] | 21[223] | GGTTATATCTTCTGTAAATCGTCGTTACATTT |
| 117 | 20[79] | 18[80] | GTTGATAACGTTCTAGCTGATAACTGAGTA |
| 118 | 20[111] | 18[112] | TAAAACTAGTAGCTATTTTTGAGATTTAGAAC |
| 119 | 20[143] | 19[159] | GAGCAAACCAGGTCATTGCCTGAGAAGAGTC |
| 120 | 20[175] | 18[176] | CGATAGCTATTTATCAAAATCATAAATACCGA |
| 121 | 20[207] | 18[208] | TTAATTTTTTTTAACCTCCGGCTTCATCT |
| 122 | 20[239] | 18[240] | TAACCTTGAACTATATGTAAATGCGAGAAAAC |
| 123 | 21[64] | 23[63] | TTAATATTACGGCGGATTGACCGTCATCGTAA |
| 124 | 21[96] | 23[95] | TTGTTAAATAACAACCCGTCGGATGACGACGA |
| 125 | 21[128] | 23[127] | AACGCCATGCCAGCTTTCATCAACACTCCAGC |
| 126 | 21[160] | 22[144] | TCAATTACTCGCGCAGAGGCGAATTCTGGCC |
| 127 | 21[192] | 23[191] | AAACATCACGCCTGATTGCTTTGAATAATGGA |
| 128 | 21[224] | 23[223] | AACAATTTCCTTTTACATCGGGAGAATTATTT |
| 129 | 22[79] | 20[80] | GGGAACAATTGTTAAAATTCGCATACCCCG |
| 130 | 22[111] | 20[112] | TGAGCGAGTCAGCTCATTTTTTAAGGTAATCG |
| 131 | 22[143] | 21[159] | TTCCTGTACAAAAATAATTCGCGTATTCATT |
| 132 | 22[175] | 20[176] | TTACAAAACTGAGCAAAAGAAGATAAACATAG |
| 133 | 22[207] | 20[208] | AACGGATTAGAAAACAAAATTAACTATTAA |
| 134 | 22[239] | 20[240] | TAACAGTACATTTGAATTACCTTTTGAGTGAA |
| 135 | 23[64] | 25[63] | CCGTGCATGCTATTACGCCAGCTGTTGGGTAA |
| 136 | 23[96] | 25[95] | CAGTATCGGTTGGGAAGGGCGATCCGTTGTAA |
| 137 | 23[128] | 25[127] | CAGCTTTCAAAGCGCCATTCGCCAATGCCTGC |
| 138 | 23[160] | 24[144] | TGATTGTTGATGGCAATTCATCAATGCCGGA |
| 139 | 23[192] | 25[191] | AGGGTTAGCGGAATTATCATCATAGATAATAC |
| 140 | 23[224] | 25[223] | GCACGTAATTTTGCGGAACAAAGAACTTTACA |
| 141 | 24[79] | 22[80] | GCCTCTTCCTGCCAGTTTGAGGGTCTCCGT |
| 142 | 24[111] | 22[112] | GCGCAACTGCCTCAGGAAGATCGCATTAAATG |
| 143 | 24[143] | 23[159] | AACCAGGCCGGCACCGCTTCTGGTATAATCC |
| 144 | 24[175] | 22[176] | TATCAGATTGGATTATACTTCTGAATACCAAG |
| 145 | 24[207] | 22[208] | AGAAGGAGAACCTACCATATCAAAAACAAT |
| 146 | 24[239] | 22[240] | CATTATCAAACAGAAATAAAGAAAATATACAG |
| 147 | 25[64] | 27[63] | CGCCAGGGCATACGAGCCGGAAGCGAGCTAAC |
| 148 | 25[96] | 27[95] | AACGACGGGAAATTGTTATCCGCTCTGCCCGC |
| 149 | 25[128] | 27[127] | AGGTCGACTTCGTAATCATGGTCACCAGCTGC |
| 150 | 25[160] | 26[144] | ACTAATAGAAAATATCTTTAGGAGGGTACCG |
| 151 | 25[192] | 27[191] | ATTTGAGGCAACAGTTGAAAGGAAAAAACAGA |
| 152 | 25[224] | 27[223] | AACAATTCCCCTCAATCAATATCTCGCCTGCA |
| 153 | 26[79] | 24[80] | CCACACAATTTTCCCAGTCACGAGGTGCGG |
| 154 | 26[111] | 24[112] | TCCTGTGTCCAGTGCCAAGCTTGCTTCAGGCT |
| 155 | 26[143] | 25[159] | AGCTCGAATCTAGAGGATCCCCGCACTAACA |
| 156 | 26[175] | 24[176] | GGTTATCTATTAGAGCCGTCAATATTCCTGAT |
| 157 | 26[207] | 24[208] | TGGCAAATATTTAGAAGTATTAGAACCACC |
| 158 | 26[239] | 24[240] | ATATCAAAGACAACTCGTATTAAATTGAGTAA |
| 159 | 27[64] | 29[63] | TCACATTACACCGCCTGGCCCTGACCCCAGCA |
| 160 | 27[96] | 29[95] | TTTCCAGTCCAGTGAGACGGGCAAGTTCCGAA |
| 161 | 27[128] | 29[127] | ATTAATGACGTATTGGGCGCCAGGAAAGAATA |
| 162 | 27[160] | 28[144] | CCGAACGACCTAAAACATCGCCATGGGAGAG |
| 163 | 27[192] | 29[191] | GGTGAGGCGGCTATTAGTCTTTAATCAATCGT |
| 164 | 27[224] | 29[223] | ACAGTGCCAGAATACGTGGCACAGCAGATTCA |
| 165 | 28[79] | 26[80] | TTGCCCTTATTGCGTTGCGCTCACACAATT |
| 166 | 28[111] | 26[112] | TCTTTTCACGGGAAACCTGTCGTGTAGCTGTT |
| 167 | 28[143] | 27[159] | GCGGTTTGATCGGCCAACGCGCGTAAAAATA |
| 168 | 28[175] | 26[176] | CTGATAGCACCACCAGCAGAAGATTTGAGGAA |
| 169 | 28[207] | 26[208] | TTTTGAATGGTCAGTATTAACACGGTCAGT |
| 170 | 28[239] | 26[240] | AAAGCGTAACGCTGAGAGCCAGCAAACCTCAA |
| 171 | 29[64] | 31[63] | GGCGAAAAATGGCCCACTACGTGAGGTGCCGT |
| 172 | 29[96] | 31[95] | ATCGGCAAGTCAAAGGGCGAAAAAAGGGAGCC |
| 173 | 29[128] | 31[127] | GCCCGAGAAGAGTCCACTATTAAAAGCCGGCG |

| | | | |
|---|---|---|---|
| 174 | 29[160] | 30[144] | ATGGAAATGCCATTGCAACAGGAATTCCAGT |
| 175 | 29[192] | 31[191] | CTGAAATGTGCTGGTAATATCCAGGAATCCTG |
| 176 | 29[224] | 31[223] | CCAGTCACGCCTGAGTAGAAGAACGGCCACCG |
| 177 | 30[79] | 28[80] | TCAGGGCGTCCTGTTTGATGGTGCAGCTGA |
| 178 | 30[111] | 28[112] | ACTCCAACAATCCCTTATAAATCAGTGGTTTT |
| 179 | 30[143] | 29[159] | TTGGAACATAGGGTTGAGTGTTGAAACGCTC |
| 180 | 30[175] | 28[176] | TACCGCCAACCTACATTTTGACGCTGCGCGAA |
| 181 | 30[207] | 28[208] | ATCGGCCTGATTATTTACATTGGACAATAT |
| 182 | 30[239] | 28[240] | CATCACTTACGACCAGTAATAAAACTGACCTG |
| 183 | 31[64] | 30[80] | AAAGCACTAAATCGGAACCCTAACCGTCTA |
| 184 | 31[96] | 30[112] | CCCGATTTAGAGCTTGACGGGGAAGAACGTGG |
| 185 | 31[128] | 31[159] | AACGTGGCGAGAAAGGAAGGGAATTAAAGGG |
| 186 | 31[160] | 30[176] | ATTTTAGACAGGAACGGTACGCCAAACAATAT |
| 187 | 31[192] | 30[208] | AGAAGTGTTTTTATAATCAGTGATCAAACT |
| 188 | 31[224] | 30[240] | AGTAAAAGAGTCTGTCCATCACGCAGTAATAA |
| 189 | 12[143] | 11[159] | AGCGTCCATAGTAAAATGTTTAGTTTATCCC |
| 190 | 9[160] | 10[144] | TCTTACCGGAAACAATGAAATAGCACTAACG |
| 191 | 7[192] | 9[191] | AGACAAAAGCAACATATAAAAGAAAAGTAAGC |
| 192 | 19[192] | 21[191] | AGACTACCCCCTTAGAATCCTTGAGATGAAAC |

### Supplementary notes

##### Thermodynamic derivation to rationalize decrease in fluorescent intensity

To explain the observed decrease in fluorescent intensity in relation to the observed $K_d$ we derived an equation to describe the fluorescence intensity as a function of receptor number and concentration of antibody. The fluorescence intensity observed in flow cytometry $F([C])$ for a given concentration $[C]$ of anti-PD1 in solution is governed by

$$F([C]) = F(0) + [F(\infty) - F(0)]\theta \qquad (1)$$

where $\theta$ is the fraction of PD1 receptors in the system bound to anti-PD1, $F(0)$ is the signal at zero concentration, and $F(\infty)$ is the signal in the limit of very large concentration. The quantity $\theta$ is given by the Langmuir adsorption isotherm

$$\theta = \frac{[C]}{K_d + [C]} \qquad (2)$$

The quantity $K_d$ is the thermodynamic dissociation constant (i.e. the reciprocal of the equilibrium constant), which relates to the standard Gibbs free energy of anti-PD1/PD1 binding via

$$\Delta G^{\circ} = RT \ln\left(\frac{K_d}{[C]^{\circ}}\right) \qquad (3)$$

(The reference concentration $[C]^{\circ}$ is the standard molar concentration of 1M.)

The fluorescence signal at very high anti-PD1 concentration is directly proportional to the total number of receptors $N_{R,tot}$ that the anti-PD1 can bind to in the whole sample, times the signal $F_1$ emitted by each bound anti-PD1 construct:

$$F(\infty) = F_1 N_{R,tot} \qquad (4)$$

Thus, plugging Eq. 2 into Eq. 1, and for mathematical clarity assuming that $F(0) = 0$, we arrive at

$$F([C]) = F_1 N_{R,tot}\left(\frac{[C]}{K_d + [C]}\right) \qquad (5)$$

This simple equation highlights how the binding fluorescence intensity depends on two key factors. The $K_d$ for binding encapsulates all of the *thermodynamic* contributions to anti-PD1/PD1 binding, e.g. enthalpy of bond formation, internal entropy change to either PD1 or anti-PD1 upon bond formation, and any non-specific additional interactions between the anti-PD1 and the cell surface below the PD1. In multivalent binding, $K_d$ is additionally a non-linear function of the number of receptors per unit area on the target surface. The $K_d$ of the anti-PD1 affects the *sharpness* of the adsorption profile measured by fluorescence, as a function of $[C]$.

The maximum fluorescence intensity $F(\infty) = F_1 N_{R,tot}$ is, on the other hand, governed by the total number of possible anti-PD1/PD1 pairs in the system, in the regime where $[C]$ is large enough that a further increase in $[C]$ leads to no increase in the fluorescence signal. It is therefore a measure of how many receptors in the sample are accessible for binding to anti-PD1. $F(\infty)$ sets the asymptotic fluorescence intensity value for the adsorption profile of the anti-PD1 at large concentration $[C]$.

The value $F(\infty)$ is controlled not only by the total number of PD1 receptors on the cell surfaces, but also how (sterically) accessible they are to the anti-PD1 construct. Large anti-PD1 constructs may be unable to access particular receptors if they are sterically blocked by other entities on the cell membrane, or if the receptor is in an undulation on the cell membrane that is too narrow for the anti-PD1 to fit into.

Two anti-PD1 constructs, one large and one small but both having an identical $K_d$, may therefore exhibit very different binding efficacies, as the larger construct may be unable to access a large fraction of the receptors in the sample due to sterics. There is also the possibility that an anti-PD1 construct with a larger $K_d$ may actually exhibit a *lower* amount of binding with the target receptors, compared to a reference anti-PD1 construct, simply because there are fewer sterically-available receptors that it can bind to on the cell surface.

A couple of practical examples of this are illustrated in **Supplementary Figure 23**. In panel **(a)**, we show a reference scenario in which we display a cell surface in green with receptors indicated as dark-green dots that can all bind free anti-PD1 antibodies with a defined $K_d$. Panel **(b)** then presents four possible scenarios for how differences in thermodynamics and sterics can affect anti-PD1 binding. For each example, a schematic plot of measured fluorescence intensity of the sample vs. anti-PD1 construct concentration is shown. Through Equation 5, the fluorescence intensity is directly proportional to the total number of receptors bound to an anti-PD1 construct.

In (i), adsorption profiles are shown for free anti-PD1, as well as for the anti-PD1-functionalized nanorods (aPD1-NR), having a larger $K_d$ than the former (indicated with orange dots). Both constructs are assumed to have the same steric accessibility to all receptors on the surface. The result is that, for a specific concentration anti-PD1, the number of surface receptors occupied by anti-PD1 is larger for the standard construct, compared to the modified construct.

In (ii), both constructs now have identical $K_d$, but aPD1-NR faces steric barriers preventing it from binding to a selection of receptors on the surface (shown as red dots). Thus, even though both have the same $K_d$, the free aPD1 exhibits more effective binding to the target surface than the larger construct.

In (iii), we show a possible outcome when aPD1-NR has a larger $K_d$ than free anti-PD1, decreasing the fraction of bound receptors $\theta$, but is also sterically blocked from binding to some of the surface receptors.

Finally in (iiii), we show a possible outcome when the larger construct has a smaller $K_d$ (shown as cyan dots) than the standard anti-PD1, but at the same time is sterically blocked from binding to a fraction of the surface receptors. This results in the interesting scenario that incubating receptors with a concentration of anti-PD1 that is similar to $K_{d,free\ anti-PD1}$ results in a higher fractional occupancy for aPD1-NR compared to free anti-PD1, while at the same time the absolute number of receptors bound by aPD1-NR is lower with respect to free anti-PD1.

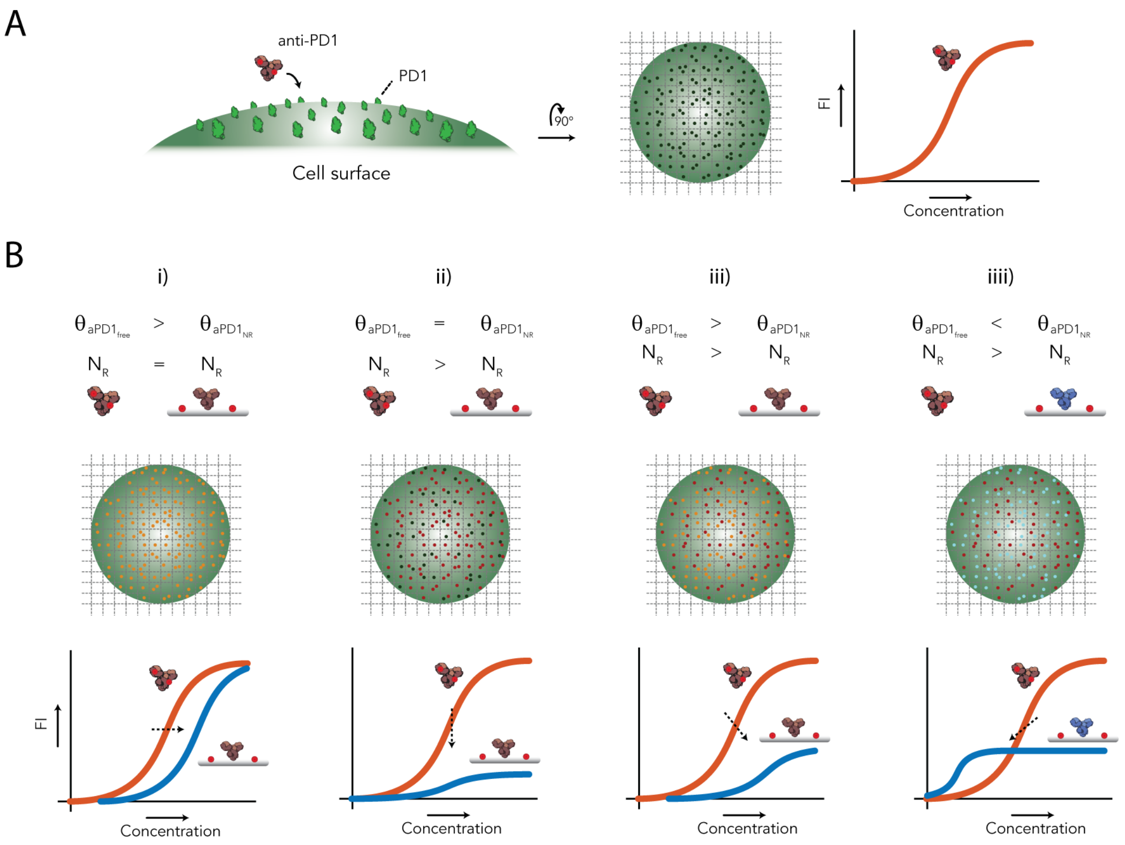

**Supplementary Figure 26 | Schematic overview of possible scenarios for anti-PD1 binding curves. (a)** The cell surface is shown in green which is decorated with PD1 receptors (dark green) that can bind to anti-PD1. A standard dose response curve for anti-PD1 binding is shown as a reference curve that indicates the relation of fluorescent intensity and concentration anti-PD1 **(b)** Four hypothetical scenarios illustrate how parameters defined in **Equation 2** and **5** impact the reference dose-response curve as shown in **(a)**. Orange dots indicate receptors that have a higher affinity to free anti-PD1 compared to aPD1-NR which results in a lower fractional occupancy $\theta$ at a fixed concentration of anti-PD1. Red dots indicate receptors that are available to free anti-PD1 but unavailable to aPD1-NR resulting in a decrease in fluorescent intensity. Cyan dots indicate receptors that have a lower affinity to free anti-PD1 compared to aPD1-NR which results in a high fractional occupancy $\theta$ at a fixed concentration of anti-PD1.
